# Neural competition predicts speech production advantages in congenital amusia

**DOI:** 10.64898/2026.11.19.767958

**Authors:** Yuchen Wang, Min Liu, Zhishuai Jin, Sizhu Huyang, Nan Guo, Jingye Zhan, Qiaoping Lian, Chuanshi Su, Qi Liu, Shuo Huang, Yunce Ji, Daxing Wu

## Abstract

Mental health research has traditionally been deficit-focused, and cognitive advantages in mental conditions have almost always been documented retrospectively. Here we propose and test an a priori pathway for predicting such advantages from a condition’s impaired neural basis, neural competition and brain–mind associations. Using congenital amusia (CA), a neurodevelopmental disorder with a well-defined deficit in right frontotemporal circuit (FTC), as a model, we predicted that interhemispheric competition would release the left homologue from inhibition, conferring advantages in speech production. Two independent studies, multimodal neuroimaging and behavioral testing, confirmed this prediction: MRI revealed right FTC impairment alongside competitive-like benefits of the left FTC in CA; and CA showed speech production advantages with intact episodic memory and constrained music learning. These findings provide the first empirical demonstration that cognitive advantages can be predicted in advance and then verified, offering a principled, mechanism-based alternative to the deficit-focused paradigm.

---

Mental health research has traditionally operated under a deficit-focused paradigm, wherein deviations from typical psychological and behavioral patterns are conceptualized as impairments requiring remediation. While this paradigm has advanced our understanding of mental disorders, it has also engendered significant limitations, including stigmatization, interpretive biases and over-pathologization^1–4^. Accumulating evidence challenges this deficit-focused paradigm and reveals that individuals with various mental conditions often exhibit some cognitive advantages^5,6^, such as enhanced creativity in bipolar disorder and dyslexia^7–10^ or superior local visual processing in autism^11^. These observations resonate with ancient wisdom—“Misfortune may be a blessing in disguise”—yet an empirical pathway to predict such advantages remains elusive.

The neural resources of the human brain are limited, and are shared and competed over by different neural systems^12–14^. In this context, among neural systems engaged in competition, if one system is inefficiency due to structural or functional impairment— that is, becomes a “weak competitor”—then the systems with which it competes may acquire greater neural resources and/or enhanced neural reserve. Neural reserve, in turn, is a key factor that positively predicts cognitive ability^15–18^. Increasing studies have confirmed the existence of neural competition, and reveal that the cognitive functions supported by neural circuits that benefit from competition are enhanced^19–27^. Functional and/or structural abnormalities in the corresponding neural systems underlie mental conditions. Therefore, like the theoretical framework proposed by Colzato et al.^28^, we argue that by investigating the neural basis of mental conditions to identify critical neural components characterized by inefficiency, inferring the neural components that may benefit from competition based on the manifestations of neural competition, and further understanding the cognitive functions supported by these potentially advantaged neural components through knowledge of brain–mind associations, it becomes possible to predict the potential cognitive advantages of people with mental conditions. However, the validity of this approach remains to be empirically tested.

Congenital amusia (CA)—a neurodevelopmental disorder characterized by impaired music processing despite normal hearing, intelligence, and musical exposure^29^—provides an ideal model for testing this framework due to its well-established neural basis and relatively low heterogeneity. The core neural basis of CA is characterized by inefficient right frontotemporal circuit (FTC)^30^, reflected in structural atrophy of the right inferior frontal gyrus (IFG)^31^, diminished top-down regulation from the right IFG to the superior temporal gyrus (STG)^32–34^, as well as reduced white matter integrity and volume of the right arcuate fasciculus (AF)^35–37^. Based on the neural basis of CA and the manifestations of neural competition^23,38,39^, we could make two inferences. First, from the perspective of interhemispheric competition between homologous circuits, the left FTC circuit and cognitive functions it supports may gain benefits. Second, from the perspective of neural competition among circuits sharing common components^24,25,40^, given the reduced modulation from the right IFG to STG, other neural circuits that rely on neural resources of the right IFG may also benefit. However, the potential presence of impaired face recognition^41,42^ and spatial processing^43–45^, both of which involve neural substrates correlated with the right IFG^46,47^, suggests that the overall neural efficiency of the right IFG may be constrained in CA. Consequently, other circuits dependent on capacity of right IFG may not exhibit competitive gains. This scenario, in turn, further highlights the potential benefit to the left frontotemporal circuit, particularly to the left IFG. Therefore, we hypothesize that CA is associated with neuroimaging features indicative of competitive benefit in the left FTC, and that speech production linked to this circuit^48–50^—particularly those related to the left IFG—represents potential cognitive advantages of CA.

Here we conducted two complementary studies to test this hypothesis. Study 1 employed multi-modal magnetic resonance imaging (MRI) to examine whether CA exhibit competitive-like enhanced neuroimaging features in left FTC. Study 2 examine whether CA exhibit advantages in speech production via storytelling task. We also accessed behavioral performance of CA in episodic memory and music processing in Study 2. The former has little relevance to the psychological and neural mechanisms of CA, and contribute to support that the speech production advantage results from specific competition rather than generalized enhancement. The latter helps to demonstrate that impairment-focused interventions may be ineffective, further highlighting the importance of recognizing and utilizing advantages. All tasks were embedded within short-term Testing-the-Limits (TtL) paradigm^51^, enabling assessment of both achievement-level (single time point) and developmental-level (developmental trajectory) cognitive advantage (See Methods).

## Results

### Study 1: Neuroimaging Features of CA in FTC

Individuals with a Montreal Battery of Evaluation of Amusia^52^ (MBEA; see Methods) score of 21.5 or lower were considered as CA^53^. According to inclusion and exclusion criteria, we collected resting-state functional, structural and diffusion MRI from 69 CA (32 males, 19.51 ± 1.90 years) and 97 controls (32 males, 19.93 ± 1.89 years). Basic characteristics for both groups are summarized in Extended Data Table. 1. To account for potential neural heterogeneity, we complemented traditional mean-based group-level analyses with a normative modeling approach^54,55^ (see Methods), which contributes to identify individual exhibiting deviate neuroimaging features from the normal range established using a large, independent Chinese reference cohort (n = 1023; see Supplementary Method 1).

**Structural MRI.** Neither group mean-based analyses nor between-group comparisons based on the proportion of deviate subjects revealed significant group differences in grey matter volume (GMV) or cortical thickness in FTC. However, a lateralization index (LI) analysis revealed that the GMV of dorsal area 44 of IFG (IFG.A44d) was significantly left-lateralized in the CA compared to controls (Fig.1a, Extended Data Table. 2; t = 3.088, *p* = 0.002, Cohen’s d = 0.497). This effect was driven by a pattern of interhemispheric competition-like alteration: CA showed numerically smaller GMV in the right IFG.A44d and larger GMV in the left IFG.A44d relative to controls, though neither was significant alone (Extended Data Fig.1a). Correlation analysis revealed that LI of IFG.A44d’s GMV was negatively associated with MBEA scores (r = −0.216, *p* = 0.007; Fig.2a). Normative model-based analyses were not conducted for LI because no acceptable model could be established.

**Resting-State functional MRI**. We examined both static and dynamic functional connectivity (sFC and dFC, respectively), as well as both static and dynamic amplitude of low-frequency fluctuations (sALFF and dALFF, respectively). Group-mean comparisons showed that sFC between the left IFG.A44v and two left temporal regions, area 41/42 of the superior temporal gyrus (STG.A41/42; t = 2.988, *p* = 0.003, Cohen’s d = 0.482) and the rostral posterior superior temporal sulcus (rpSTS; t = 3.063, *p* = 0.003, Cohen’s d = 0.494), was significantly higher in CA than in controls (Fig.1b, Extended Data Table. 2). Normative modeling analysis (Table. 1), using a deviation threshold of z = 0, identified a greater proportion of CA subjects with positive deviations in sFC between left IFG.A44v and left STG.A41/42 (66.15% in CA vs 41.49% in controls, *p* = 0.002), rpSTS (67.69% in CA vs 43.61% in controls, *p* = 0.003), and caudal area 22 of STG (STG.A22c; 63.08% in CA vs 37.23% in controls, *p* = 0.001). However, no significant between-group differences emerged when applying the more stringent threshold of |z| = 1.96. Correlation analyses revealed that both raw sFC and its deviation scores between left IFG.A44v and rpSTS were negatively correlated with MBEA scores (r = −0.180, *p* = 0.024 for raw sFC, Fig.2b; r = −0.172, *p* = 0.030 for deviation scores, Fig.3a). No significant group differences were found for sFC’s LI in FTC under either analytical approach.

**Table. 1.** Results of Individual Deviation Levels of MRI Metrics.

| Metrics and Region | RMSE | SMSE | CA | Controls | χ<sup>2</sup> | *p* |
|---|---|---|---|---|---|---|
| sFC (z = 0) |  |  |  |  |  |  |
| L_IFG.A44v – L_STG.A41/42 | 0.220 | 0.969 | 43/22 | 39/55 | 9.360 | 0.002 |
| L_IFG.A44v – L_STG.A22c | 0.211 | 0.999 | 44/21 | 41/53 | 2.988 | 0.004 |
| L_IFG.A44v – L_rpSTS | 0.224 | 0.993 | 41/24 | 35/59 | 3.063 | 0.001 |
| dALFF (z = ±1.96) |  |  |  |  |  |  |
| L_IFG.A45c (30-volume) | 0.070 | 0.490 | 6/0 | 0/0 | 9.017 | 0.003 |
| L_IFG.A44op (30-volume) | 0.075 | 0.838 | 6/0 | 0/0 | 9.017 | 0.003 |
| L_IFG.A44v (30-volume) | 0.101 | 0.758 | 6/0 | 0/0 | 9.017 | 0.003 |
| dFC |  |  |  |  |  |  |
| R_IFG.IFS – R.STG.A22c<br>(30-volume, z = ±1.96) | 0.036 | 0.986 | 5/0 | 0/0 | 7.466 | 0.006 |
| L_IFG.A44v – L_STG.A41/42<br>(30-volume, z = 0) | 0.030 | 1.008 | 19/46 | 50/44 | 8.981 | 0.003 |
| L_IFG.A44v – L_STG.A22c<br>(30-volume, z = 0) | 0.035 | 0.999 | 19/46 | 48/46 | 7.513 | 0.006 |
| dFC-LI (z = 0) |  |  |  |  |  |  |
| IFG.A44v – STG.A38m<br>(30-volume) | 0.293 | 0.977 | 26/39 | 59/35 | 8.005 | 0.005 |
| FA of R.dAF (z = 0) | 0.016 | 0.703 | 24/45 | 54/41 | 7.799 | 0.005 |
| RD of R.dAF<sup>a</sup> (z = 0) | 0.203 | 0.715 | 47/22 | 38/57 | 12.655 | 0.004 |
| FA-LI of dAF (z = 0) | 0.015 | 0.911 | 42/27 | 37/58 | 7.694 | 0.006 |
| Volume-LI of dAF (z = 0) | 0.110 | 0.998 | 22/47 | 51/44 | 7.691 | 0.006 |

*Note:* For each group, deviations are presented as the number of individuals with positive deviations / the number with negative deviations (e.g., 6/0 indicates 6 individuals with positive deviations and 0 with negative deviations in that group). z = 0 indicates that individuals were identified as deviating using 0 as the threshold, and z = ±1.96 indicates identification using ±1.96 as the threshold. RMSE and SMSE represent the fitting errors of the corresponding normative models for each metric. Superscript a: the actual value of RMSE is 10<sup>-4</sup> times the displayed value. CA, congenital amusia; sFC, static functional connectivity; dFC, dynamic functional connectivity; FA, fractional anisotropy; RD, radial diffusivity; dALFF, dynamic amplitude of low-frequency fluctuations; LI, lateralization index; IFG, inferior frontal gyrus; STG, superior temporal gyrus; A44v, ventral area 44; A45c, caudal area 45; A44op, opercular area 44; A38m, medial area 38; A22c, caudal area 22; IFS, inferior frontal sulcus; rpSTS, rostral posterior superior temporal sulcus; dAF, direct arcuate fasciculus; L, left; R, right; RMSE, root mean square error; SMSE, standardized mean square error.

For dFC, computed using a sliding-window approach, analyses were performed at window sizes of 30 and 50 volumes. The 30-volume window served as the primary analysis, and results at 50 volumes were used to assess robustness. Unless otherwise specified, results reported below are based on the 30-volume window, and findings that also held at 50-volume window are explicitly note. Group-mean comparisons revealed significantly lower LI between IFG.A44v and multiple STG subregions in CA compared with controls (all *p*s < 0.01, reproduced at window sizes of 30 and 50; Fig.1e-f, Extended Data Table. 2). These effects were also driven by an interhemispheric competition-like alteration, with CA showing numerically reduced dFC in the left hemisphere and increased dFC in the right hemisphere, though neither alone was significant (Extended Data Fig.1c-d). Normative modeling analysis with a z = 0 threshold (Table. 1) indicated a higher proportion of CA subjects with negative deviations in dFC between left IFG.A44v and left STG.A41/42 (70.77% in CA vs 46.81% in controls, *p* = 0.003) and STG.A22c (70.77% in CA vs 48.94% in controls, *p* = 0.006), as well as negative deviations in LI of dFC between IFG.A44v and medial area 38 of STG (STG.A38m; 60.00% in CA vs 37.23% in controls, *p* = 0.005). When using a stricter threshold of |z| = 1.96, more CA subjects exhibited positive deviations in dFC between the right inferior frontal sulcus of IFG (IFG.IFS) and right STG.A22c (7.69% in CA vs 0% in controls, *p* = 0.006; Table. 1). Correlation analyses further showed that deviation scores for dFC between right IFG.IFS and right STG.A22c were negatively correlated with MBEA scores (r = −0.176, *p* = 0.027; Fig.3c), whereas the LI of dFC between IFG.A44v and STG was positively correlated with MBEA scores (all *p*s < 0.05; all reproduced at window sizes of 30 and 50, except for the LI of dFC between IFG.A44v and STG.A41/42; Fig.2d-g); in addition, deviation scores of the LI of dFC between IFG.A44v and STG.A38m were positively correlated with MBEA scores (r = 0.286, *p* < 0.001; Fig.3d).

Like dFC, dALFF analyses were also performed at window sizes of 30 and 50 volumes to assess robustness; findings that also held at 50 volumes are explicitly noted. Group-mean comparisons showed significantly higher values in the left IFG.A44v (*p* < 0.001; reproduced at window sizes of 30 and 50) and left caudal area 45 of IFG (IFG.A45c; t = 3.199, p = 0.002, Cohen’s d = 0.516) in CA (Fig.1c, Extended Data Table. 2). Normative modeling at the |z| = 1.96 threshold revealed a greater proportion of CA subjects with positive deviations in left IFG.A44v, IFG.A45c, and the opercular area 44 of IFG (IFG.A44op; all *p*s = 0.003; Table. 1); no significant results were found when using the z = 0 threshold. All these dALFF alterations were negatively correlated with MBEA scores (all *p*s < 0.05; Fig.2c, Fig.3b).

**Table. 2.** Results of Individual Deviation Levels (Development-level) of Behavioral Metrics.

| Metrics | RMSE | SMSE | CA | Controls | χ<sup>2</sup> | *p* |
|---|---|---|---|---|---|---|
| MAI（z = 0） | 64.907 | 0.993 | 11/19 | 65/38 | 6.632 | 0.010 |
| Proportion of medium silent pause duration（z = 0） | 2.157 | 0.861 | 8/22 | 50/53 | 4.522 | 0.034 |
| Syntactic Complexity（z = 0） | 0.250 | 0.903 | 23/7 | 54/49 | 5.600 | 0.018 |
| Local Coherence（z = 0） | 0.040 | 0.750 | 21/9 | 48/55 | 5.095 | 0.024 |

*Note:* For each group, the number of individuals showing different cognitive characteristics is presented as the number with cognitive advantage / the number with cognitive disadvantage (e.g., 4/0 indicates that 4 individuals in the group showed a cognitive advantage and 0 showed a cognitive disadvantage). z = 0 indicates that individuals were classified using 0 as the threshold. RMSE and SMSE represent the fitting errors of the corresponding normative models for each metric. CA, congenital amusia; MAI, mean asynchrony index; RMSE, root mean square error; SMSE, standardized mean square error.

No significant group differences were observed for sALFF under either analytical approach.

**Diffusion MRI.** We reconstructed three sub-segments of AF using probabilistic tractography: the anterior (aAF), direct (dAF) and posterior segment (pAF). The reconstructed sub-segments were presented in the Supplementary Fig.2. We then assessed their macro– and microstructural properties. Group-mean comparisons of white matter microstructure revealed significantly lower fractional anisotropy (FA) in the right dAF (t = −2.713, *p* = 0.007, Cohen’s d = −0.430; Fig.1d) and aAF (t = –2.730, *p* = 0.005, Cohen’s d = −0.432) in CA (Fig. 1d, Extended Data Table. 2). In addition, CA showed significantly higher LI of FA in the dAF (t = 2.426, p = 0.016, Cohen’s d = 0.384; Fig.1a, Extended Data Table. 2), while FA in the left dAF was also numerically lower (Extended Data Fig.1b). Normative modeling (Table. 1) at the z = 0 threshold identified a greater proportion of CA subjects with negative deviations in FA of right dAF (65.22% in CA vs 43.16% in controls, *p* = 0.005) and LI of dAF’s volume (68.12% in CA vs 46.32% in controls, *p* = 0.006), as well as positive deviations in radial diffusivity (RD) of right dAF (68.12% in CA vs 40.00% in controls, *p* = 0.004) and LI of FA in dAF (60.87% in CA vs 38.95% in controls, *p* = 0.006); no significant differences were detected at the |z| = 1.96 threshold. Correlation analyses showed that FA of the right dAF and aAF (Fig.2h), as well as deviation scores of FA of the right dAF (Fig.3e), were positively correlated with MBEA scores; in contrast, LI of FA of the right dAF (Fig.2i) and its deviation scores (Fig.3f) were negatively correlated with MBEA scores (all *p*s < 0.05).

**Fig. 1.**
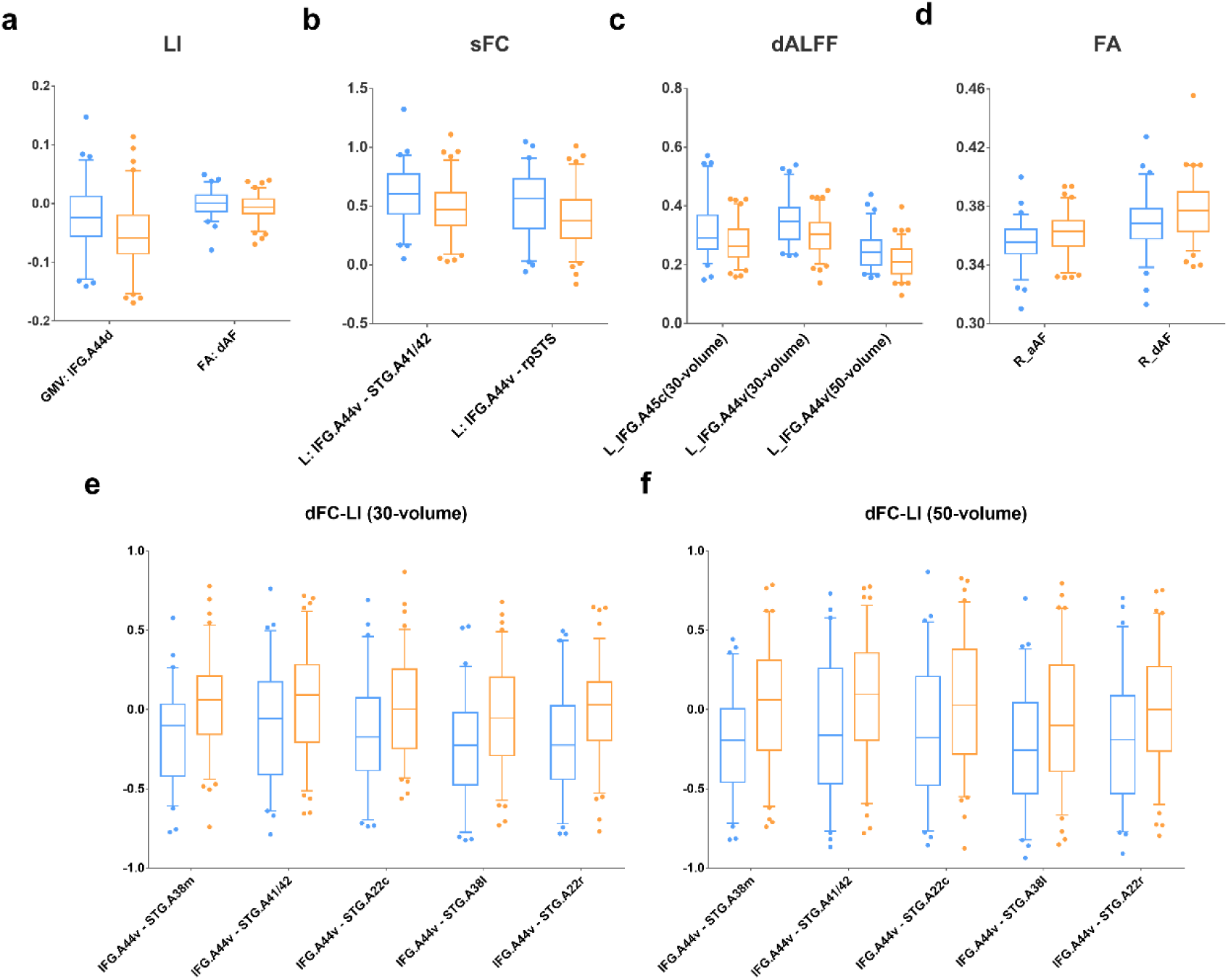
Box-plots of MRI metrics showing significant group mean difference. **a**, congenital amusia showed enhanced leftward lateralization in GMV of IFG.A44d and FA of dAF. **b,** congenital amusia showed enhanced sFC between left IFG.A44v and STG. **c,** congenital amusia showed enhanced dALFF in left IFG. **d,** congenital amusia showed decreased FA in right aAF and dAF. **e-f,** congenital amusia showed decreased leftward lateralization in dFC between IFG and multiple STG sub-regions. Orange represents controls and blue represents congenital amusia. The upper and lower bars represent 95 and 5 percentiles, respectively. LI, lateralization index; GMV, grey matter volume; sFC, static functional connectivity; dALFF, dynamic amplitude of low-frequency fluctuations; FA, fractional anisotropy; dFC, dynamic functional connectivity; IFG, inferior frontal gyrus; STG, superior temporal gyrus; A44d, dorsal area 44; A41/42, area 41/42; A44v, ventral area 44; A45c, caudal area 45; A38m, medial area 38; A22c, caudal area 22; A38l, lateral area 38; A22r, rostral area 22; rpSTS, rostral posterior superior temporal sulcus; aAF, anterior arcuate fasciculus; dAF, direct arcuate fasciculus; R, right; L, left.

**Fig. 2.**
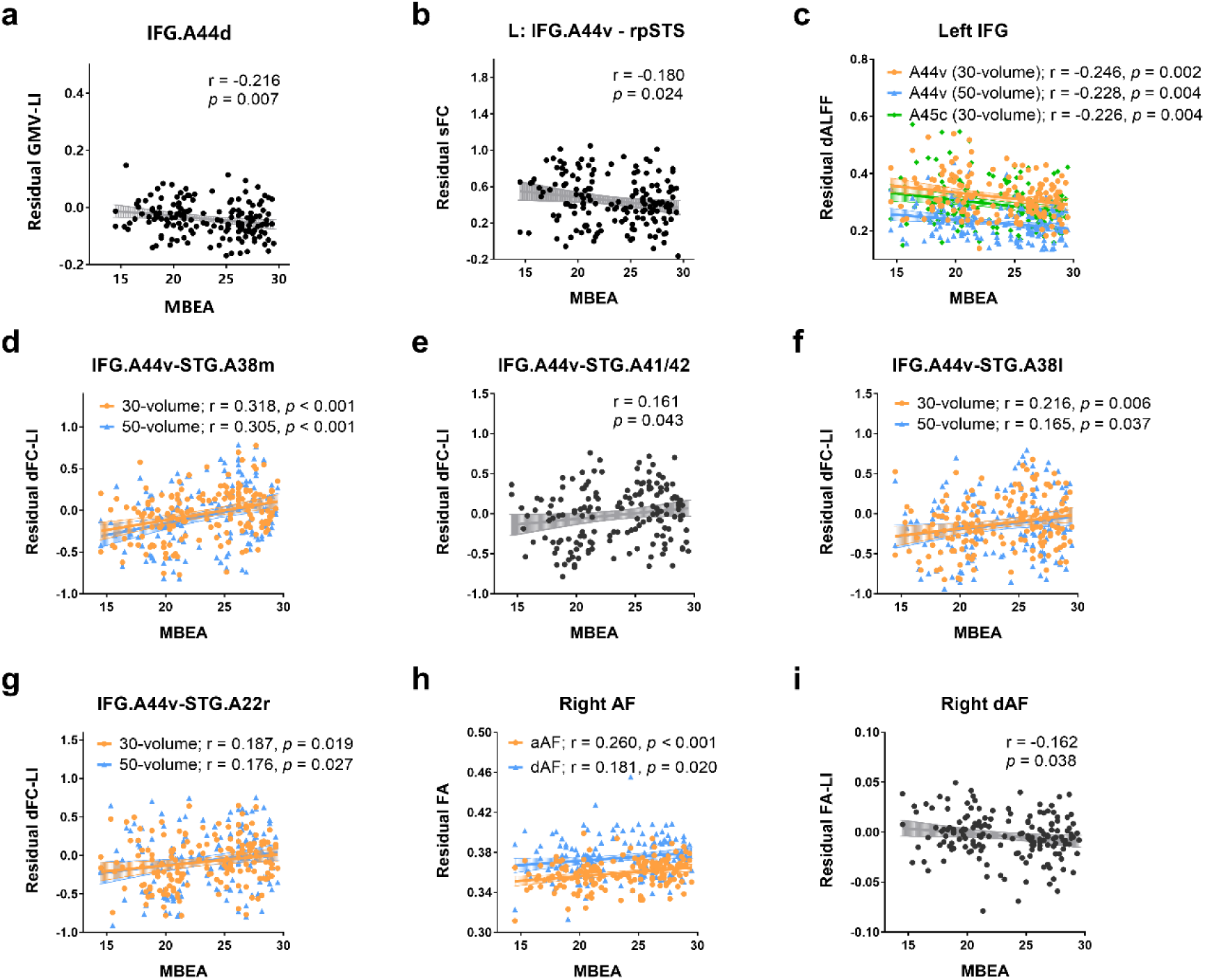
Correlation between MBEA and MRI metrics showing significant group mean difference. We regressed out the effect of sex, age and education years from all metrics; for rs-fMRI metrics, mean FD was additionally regressed out; for sMRI metrics, image quality rating and total intracranial volume were additionally regressed out. MBEA, Montreal Battery of Evaluation of Amusia; LI, lateralization index; GMV, grey matter volume; sFC, static functional connectivity; dALFF, dynamic amplitude of low-frequency fluctuations; FA, fractional anisotropy; dFC, dynamic functional connectivity; IFG, inferior frontal gyrus; STG, superior temporal gyrus; A44d, dorsal area 44; A41/42, area 41/42; A44v, ventral area 44; A45c, caudal area 45; A38m, medial area 38; A38l, lateral area 38; A22r, rostral area 22; rpSTS, rostral posterior superior temporal sulcus; AF; arcuate fasciculus; aAF, anterior AF; dAF, direct AF; L, left.

### Study 2: Behavioral Performance of CA in Speech Production, Episode Memory and Music Processing

We recruited an independent sample of 30 CA subjects and 103 healthy controls between December 2024 and October 2025, using the same screening procedures as Study 1. Both groups did not differ in sex (*p* = 0.658). The CA was slightly older (*p* = 0.033), had longer years of education (*p* = 0.012), and scored lower on the Wechsler Adult Intelligence Scale (WAIS; *p* = 0.003) than controls; these variables were controlled for in subsequent analyses when they showed significant associations with dependent measures. CA scored significantly lower on all MBEA subtests and on the Goldsmiths Musical Sophistication Index (Gold-MSI) subscales of active engagement, perceptual abilities, singing ability, and general sophistication (all *p*s < 0.05). No significant self-reported differences in language expression ability were found between groups. More detailed information for both groups are summarized in Extended Data Table. 3.

Because group-mean comparisons may be insensitive to advantages carried by only a minority of individuals and paradigms we used here lack standard population norms, we additionally applied a normative modelling approach similar to that used in Study 1 to identify individuals with potential cognitive advantage (see Methods). Briefly, for achievement-level normative modelling, models were built from controls’ pre-test data, and leave-one-out cross-validation (LOOCV) was used to evaluate the models and generate deviation scores for controls’ pre-test data; acceptable models were then applied to controls’ post-test data and each stage’s data of CA to compute deviation scores. Similarly, for developmental-level normative modelling, pre-test scores were added as predictors to build normative models for post-test from controls’ data, and LOOCV was used to evaluate the models and generate deviation scores; acceptable models were then applied to CA to compute deviation scores, which quantify training gains relative to each individual’s baseline and demographic profile, thereby identifying potential cognitive advantage on the developmental dimension.

**Speech production performance.** Our storytelling task used a picture sequence with 30 events, divided into seven episodes that together formed a single narrative, to elicit connected speech. Participants were asked to view the pictures for 5 minutes and then narrated the story with no time limit; between the pre-test and post-test narrations, they received a brief, structured narrative strategy guidance. Fluency, micro– and macro-linguistic measures were used to assess participants’ performance. For more detailed information about the task and measurements, see Methods.

Mixed linear model (MLM) analysis revealed that, after receiving narrative strategy instruction, both groups produced more total number of words (TNW; *F*_(1, 131.000)_ = 163.935, *p* < 0.001) and total number of utterances (TNU; *F*_(1, 131.000)_ = 117.825, *p* < 0.001); both groups also exhibited greater lexical diversity (*F*_(1, 131.000)_ = 54.731, *p* < 0.001), global coherence (*F*_(1, 262)_ = 23.944*p* < 0.001), and theme-search performance (*F*_(1, 131.000)_ = 35.993, *p* < 0.001), and accurately recounted more events (*F*_(1, 131.000)_ = 33.210, *p* < 0.001) while omitting fewer (*F*_(1, 131.000)_ = 7.938, *p* = 0.006). At the same time, however, both groups showed increased total number of silent pauses frequency (*F*_(1, 131.000)_ = 6.870, *p* = 0.010) and overall pause frequency (*F*_(1, 131.000)_ = 4.522, *p* = 0.035), a shorter mean length of utterance (MLU; *F*_(1, 131.000)_ = 8.254, *p* = 0.005), and decreased local coherence (*F*_(1, 131.000)_ = 11.183, *p* = 0.001) and informativeness density/efficiency (*F*_(1, 131.000)_ = 24.299, *p* < 0.001; *F*_(1, 131.000)_ = 15.418, *p* < 0.001). The above findings are shown in Fig. 4. In addition, CA exhibited several characteristics distinct from controls, including a faster speech rate (*F*_(1, 131.000)_ = 6.848, *p* = 0.010), a higher frequency (*F*_(1, 131.000)_ = 4.060, *p* = 0.046) and proportion of duration (*F*_(1, 131.000)_ = 4.943, *p* = 0.028) of short silent pauses, more voiced pauses (*F*_(1, 129.338)_ = 7.730, *p* = 0.006; specifically filled pauses, *F*_(1, 129.373)_ = 9.296, *p* = 0.003), a lower frequency (*F*_(1, 131.000)_ = 5.009, *p* = 0.027) and proportion of duration (*F*_(1, 131.000)_ = 5.015, *p* = 0.027) of long silent pauses, a longer MLU (*F*_(1, 131)_ = 4.831, *p* = 0.030), a higher frequency of total pauses (*F*_(1, 130.998)_ = 7.914, *p* = 0.006) and higher informativeness efficiency (*F*_(1, 131.000)_ = 4.981, *p* = 0.027). Furthermore, MLM analysis also revealed a significant Group × Time interaction in syntactic complexity (*F* _(1, 131.000)_ = 4.602, *p* = 0.034); simple effect analysis showed that syntactic complexity in the post-test was comparable to that in the pre-test in CA (*p* = 0.839), whereas syntactic complexity in controls decreased at the post-test stage (*p* < 0.001). Above findings were presented in Fig.4. The above findings are presented in Fig.4h, Fig.4i, Fig.4l, Fig.5 and Extended Data Table. 4-6.

**Fig. 3.**
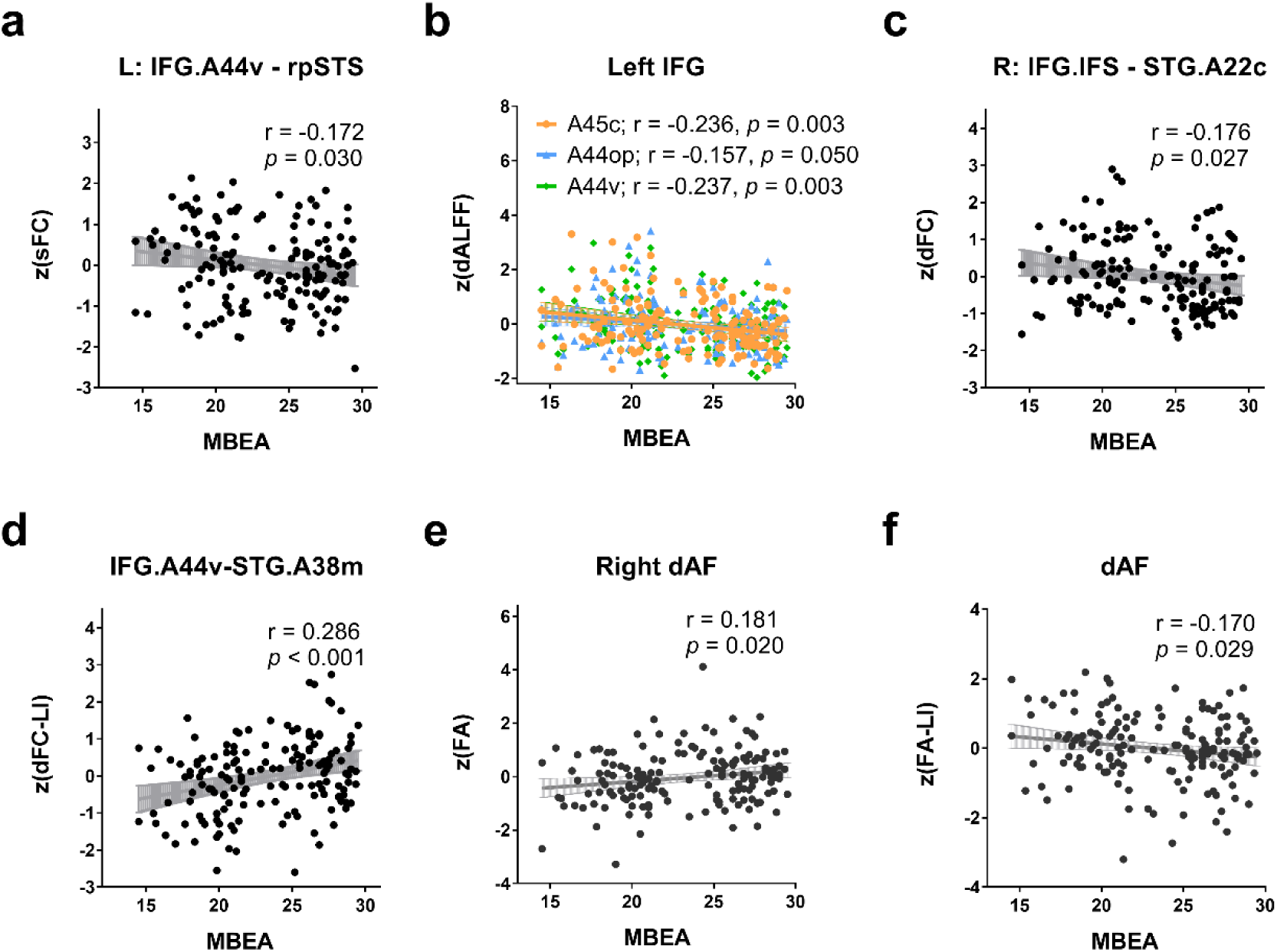
Correlation between MBEA and deviation score of MRI metrics showing significant group difference. MBEA, Montreal Battery of Evaluation of Amusia; LI, lateralization index; sFC, static functional connectivity; dALFF, dynamic amplitude of low-frequency fluctuations; FA, fractional anisotropy; dFC, dynamic functional connectivity; IFG, inferior frontal gyrus; STG, superior temporal gyrus; A44v, ventral area 44; A45c, caudal area 45; A44op, opercular area 44; A38m, medial area 38; A22c, caudal area 22; IFS, inferior frontal sulcus; rpSTS, rostral posterior superior temporal sulcus; dAF, direct arcuate fasciculus; L, left; R, right.

**Fig. 4.**
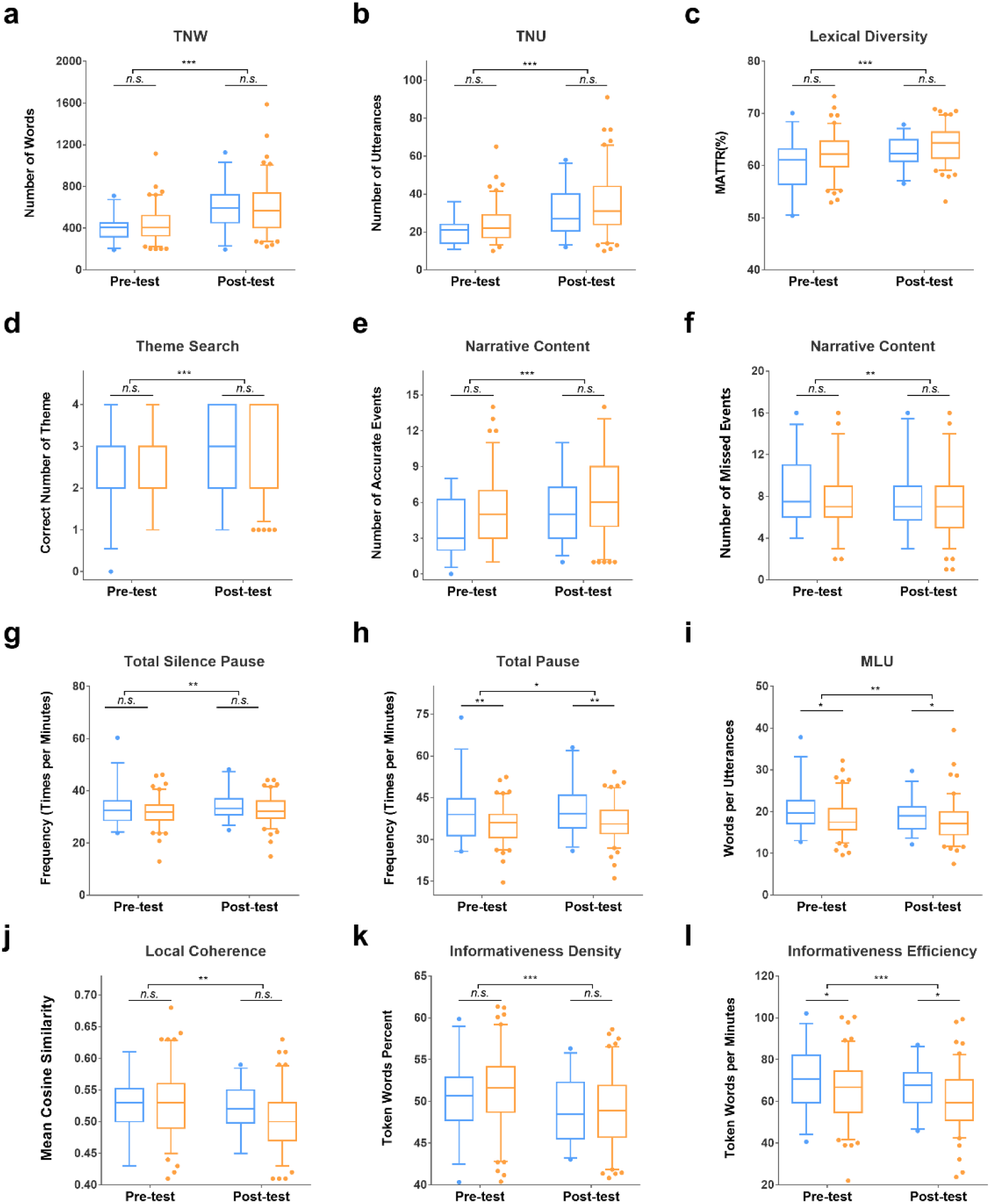
Box-plots of metrics showing significant time-effect in speech production tasks. **a-e**, performance during the post-test were higher than the pre-test. **f,** both groups missed few events during the post-test than the pre-test. g-h, both groups showed higher frequency of pause during the post-test than the pre-test; congenital amusia also showed higher frequency of total pause than controls during the pre– and post-test (h). **i,** both groups showed longer MLU during the post-test than the pre-test; congenital amusia also showed longer MLU than controls during the pre– and post-test. **j-k**, both groups showed lower post-test than pre-test performance. **l,** both groups showed lower efficiency during the post-test than the pre-test; congenital amusia also showed higher efficiency than controls during the pre– and post-test. In **d,** the median of congenital amusia during pre-test was 2, and the median during pretest and posttest of the control group were 2 and 4, respectively. Orange represents controls and blue represents congenital amusia. The upper and lower bars represent 95 and 5 percentiles, respectively. TNW, total number of words; TNU, total number of utterances; MATTR, moving average type-token ratio; MLU, mean length of utterance. *n.s.*, no significant; *, *p* < 0.05; **, *p* < 0.01; ***, *p* < 0.001.

**Fig. 5.**
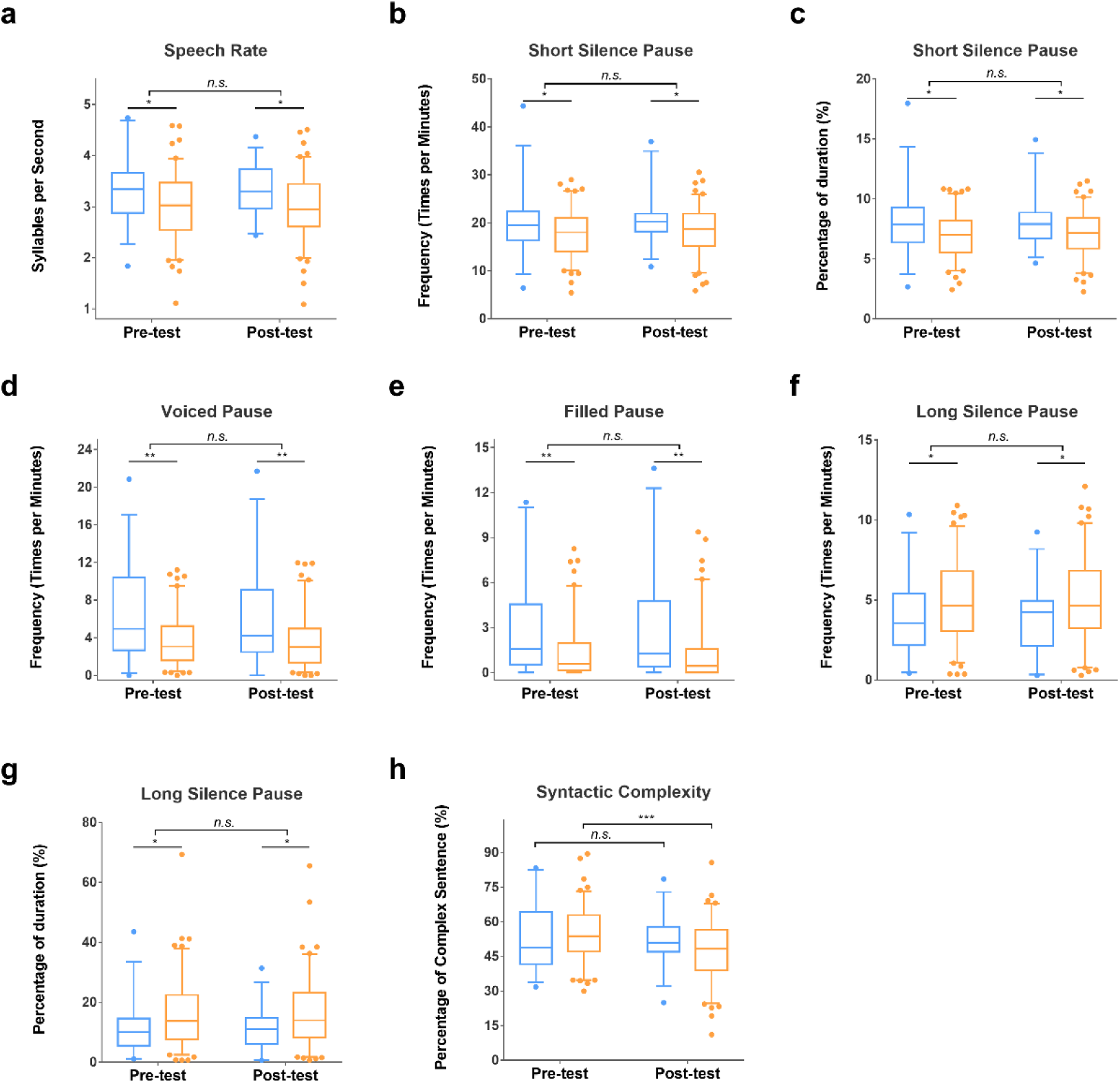
Box-plots of metrics showing significant group-specific effect in speech production tasks. **a**, congenital amusia showed faster speech rate than controls. **b-c**, congenital amusia showed higher frequency and proportion of duration of short silent pause. **d-e,** congenital amusia showed more voiced pauses, especially filled pauses than controls. **f-g,** congenital amusia showed lower frequency and proportion of duration of long silent pause. **h**, congenital amusia showed comparable syntactic complexity in the post-test and pre-test, whereas controls showed decreased syntactic complexity at the post-test stage. Group effects were also observed in total pause frequency, mean length of utterance and informativeness efficiency, which have been presented in Fig. 4 and are therefore not repeated here. *n.s.*, no significant; *, *p* < 0.05; **, *p* < 0.01; ***, *p* < 0.001

**Fig. 6.**
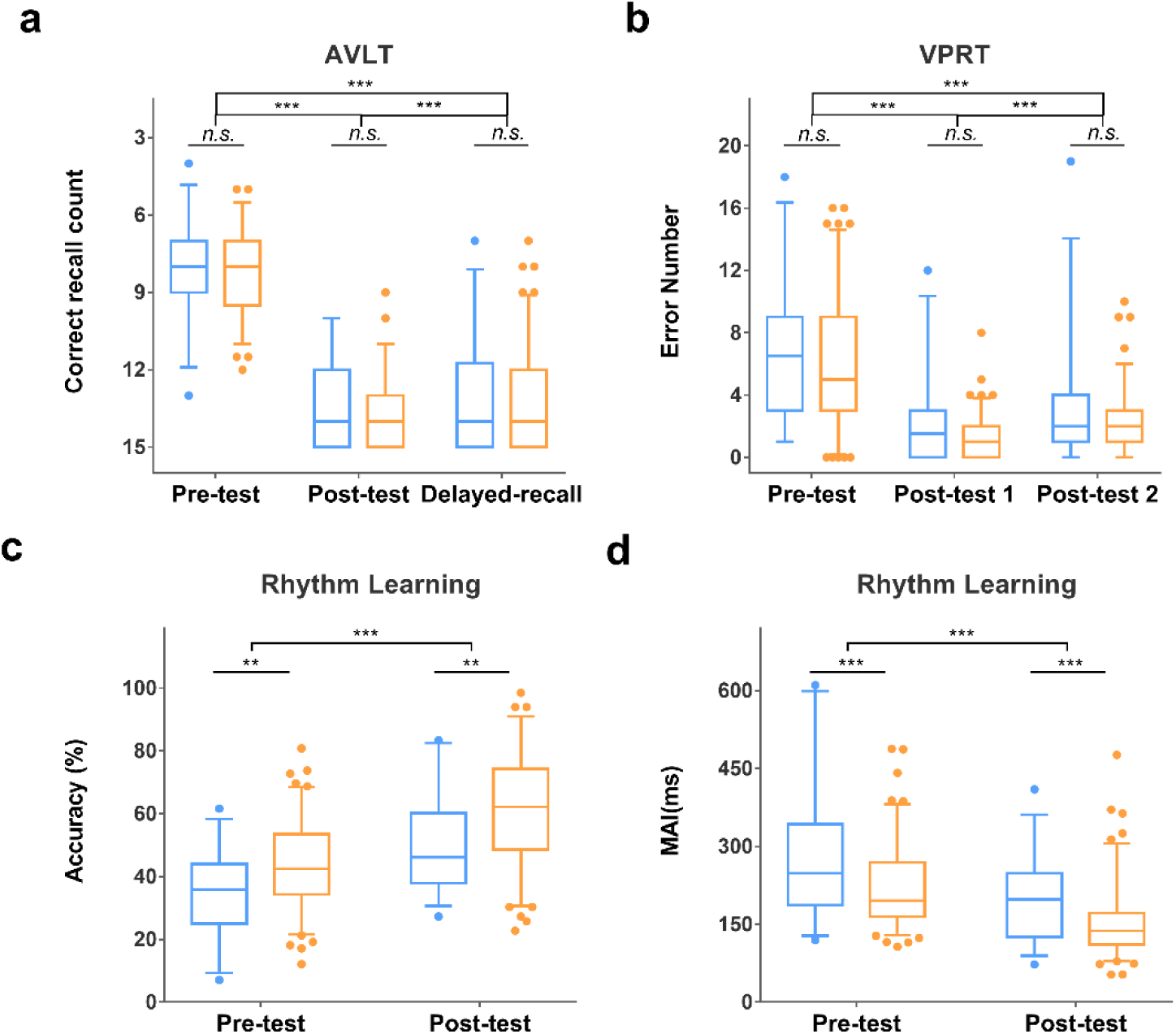
Box-plots of performance on the episodic memory and rhythm learning tasks. **a**, both groups recalled more words at the post-test and delayed-recall stages than at the pre-test; both groups also recalled more words at the post-test than at the delayed-recall stage. **b**, both groups recognized more pictures at both post-test stage than at the pre-test; both groups also recognized more pictures at the post-test 1 than at the post-test 2. **c,** both groups accurately reproduced more notes at the post-than at the pre-test; congenital amusia accurately reproduced fewer notes than controls at both pre– and post-test. **d,** both groups showed lower MAI at the post-than at the pre-test; congenital amusia showed higher MAI than controls at both pre– and post-test. MAI, mean asynchrony index; AVLT, auditory verbal learning test; VPRT, visual picture recognition test. *n.s.*, no significant; **, *p* < 0.01; ***, *p* < 0.001.

As shown in Table. 2, normative modelling at the z = 0 threshold revealed a greater proportion of CA subjects with positive deviations in local coherence (70% in CA vs 46.60% in controls; χ^2^ = 5.095, *p* = 0.024) and syntactic complexity (76.67% in CA vs 52.43% in controls; χ^2^ = 5.600, *p* = 0.018) at the developmental level, as well as negative deviations in the proportion of duration of medium silent pauses (73.33% in CA vs 51.46% in controls; χ^2^ = 4.522, *p* = 0.034) at the developmental level. Other measures either failed to yield acceptable normative models or showed no significant group differences in normative modelling analysis.

**Episodic memory performance.** Episodic memory was measured by two TtL paradigm adapted tasks: an adapted Auditory Verbal Learning Test (AVLT)^56^ and a Visual Picture Recognition Test (VPRT)^57^. For more detailed information about both tasks, see Methods.

MLM analysis revealed significant Time effects in both tasks (AVLT: *F*_(2, 109.296)_ = 90.410, *p* < 0.001; VPRT: F_(2, 163.848)_ = 628.944, *p* < 0.001), whereas neither task showed a significant Group effect or Group × Time interaction (all *p*s > 0.05). For AVLT, post-hoc analysis indicated that both groups recalled more words at the post-test and delayed-recall stages than at the pre-test stage (*p*s < 0.001); both groups also recalled more words at the post-test stage than at the delayed-recall stage (*p* < 0.001). For the VPRT, post-hoc analysis indicated that both groups successfully recognized more pictures at both post-test stage than at the pre-test stage (*p*s < 0.001); both groups also successfully recognized more pictures at the first than at the second post-test stage (*p* < 0.001). The above findings are shown in Fig.6a, Fig.6b and Extended Data Table. 7. Normative modelling analysis revealed no group differences in both tasks at the development-level; acceptable normative models were failed to yield at achievement-level.

**Music rhythm learning performance.** We administered a short-term music rhythm learning task^15,58^ to evaluate music processing ability. In a short, participants needed to learn and imitate three rhythm sequences, each composed of 11 notes with same pitch, duration, and inter-note intervals, but differing in the ordering of those intervals. The three sequences use the same notes and intervals, but the interval order differs. Task performance is assessed by response accuracy and mean asynchrony index (MAI). For more detailed information about both tasks, see Methods.

As shown in Fig.6c, Fig.6d and Extended Data Table. 7, MLM analysis revealed significant main effects of Group (Accuracy: *F*_(1, 131.291)_ = 8.018, *p* = 0.005; MAI: *F*_(1, 131)_ = 14.391, *p* < 0.001) and Time (*F*_(1, 131.000)_ = 128.686, *p* < 0.001; *F*_(1, 131.000)_ = 78.185, *p* < 0.001), but no significant Group × Time interaction (Accuracy: *F*_(1, 131.000)_ = 0.415, *p* = 0.521; MAI: *F*_(1, 131.000)_ = 1.371). This pattern of effects consistent with the core impairment of CA: although both groups improved in accuracy and MAI after training, yet CA performed worse than controls in both measures at both pre-test and post-test. Normative modelling analysis further indicated that fewer CA participants than controls showed positive deviations in MAI development (36.67% in CA vs 63.11% in controls; χ^2^ = 6.632, *p* = 0.010; Table. 2). Since lower MAI reflects better performance, this deviation pattern suggests that, despite their improvement trajectory not differing detectably at the group-mean level, CA struggled to align their response patterns more precisely with the targets. Normative modelling analysis revealed no group differences in accuracy at the development-level; acceptable normative models were failed to yield at achievement-level.

## Discussion

This study set out to test whether cognitive advantages in a mental condition can be predicted, rather than merely documented post hoc, from knowledge of the condition’s impaired neural basis combined with neural competition and brain-mind associations. Using CA as a model, we reasoned that the well-characterized inefficiency of the right FTC should, through interhemispheric competition, release the left homologue from cross-hemispheric inhibition and enable it to accrue neural reserve, and that the cognitive functions supported by the left FTC—speech production—should therefore be enhanced.

Across two independent studies, multimodal neuroimaging and behavioral evidence converged to support this prediction. Generally, Study 1 indicated that CA showed structural and functional impairments of right FTC alongside competitive-like benefits in the left FTC; Study 2 further demonstrated that CA presented advantages in speech production while exhibiting intact episodic memory and constrained music learning. These findings provide initial empirical validation for a principled, a priori pathway to predict cognitive advantages in individuals with mental conditions, moving beyond retrospective explanations of “misfortune may be a blessing in disguise.”

### Inefficient right FTC in CA

**Structural impairments in right FTC.** Study 1 found reduced FA in the right aAF and dAF, as well as a higher proportion of subjects with negative FA deviations and positive RD deviations in the right dAF in CA. FA and RD are key indices of white matter microstructural integrity, with reduced FA accompanied by elevated RD suggesting demyelination or axonal damage^59,60^. This interpretation was further confirmed by the positive correlation between right AF’s FA and MBEA scores, which links the observed microstructural abnormality to the music perception deficit of CA. These findings align with previous reports that right AF integrity predicts musical performance^15^ and that impaired right frontotemporal white matter integrity is a core feature of CA^35,36^. For gray matter, CA showed a trend toward reduced GMV of right IFG, which was the right-hemisphere component of a significant competitive-like leftward shift in GMV lateralization of (left numerically higher, right numerically lower). This is consistent with prior studies reporting right IFG atrophy in CA^31^. Together, these findings suggest structural impairments of the right FTC in CA. The present study also found some structural features of the right FTC in CA that differ from previous studies, these are discussed in the Supplementary Discussion 1.

**Unstable right frontotemporal functional communication.** Similar to GMV, CA showed a trend toward increased dFC between the right IFG and STG, which was the right-hemisphere component of a significant competitive-like rightward shift in dFC lateralization of (left numerically lower, right numerically higher). In addition, individual-level normative modelling further revealed a higher proportion of subjects with positive dFC deviations in CA. Although no unilateral dFC result survived after multiple comparison correction in the group-mean analysis, these findings consistently suggest higher dFC between right IFG and STG in CA. Previous findings indicated that damage to white matter pathways disrupts the stability of signal transmission between nodes^61–63^. The present study and earlier studies consistently found impaired white matter integrity of the right AF ^35,36^, a key pathway connecting IFG and STG, in CA. Therefore, the observed increase may reflect unstable functional coordination between the right IFG and STG in CA. Consistent with this view, the present study found that the deviation score of right frontotemporal dFC was negatively correlated with MBEA scores, and that relatively greater right-hemisphere dFC was also negatively correlated with MBEA scores. To our knowledge, this study provides the first exploration of dFC features in CA. These findings extend our understanding of the neural basis of CA, indicating that CA involves not only reduced functional connectivity strength between the right IFG and STG^32–34^, but also unstable functional communication. Such an unstable interaction may weaken the capacity of the IFG to exert sustained and stable top-down control over perceptual processing in the STG. Together, these structural and functional alterations may constitute a more complete picture of attenuated top-down modulation of CA.

**Limited learning efficiency in music.** Besides lower MBEA score, Study 2 also found CA subjects self-reported worse musical perception and singing ability (fewer engagement may be attributed to poorer music ability). Further, in the rhythm learning task, both groups improved their performance with practice, indicating preserved plasticity in CA. However, CA performed worse than controls at both pre– and post-test and fewer CA individuals showed developmental advantages in MAI. These findings suggested that, the music deficit is therefore not an all-or-none loss but a constraint on learning efficiency in CA. Mechanistically, the right IFG is critically involved in predictive coding, error monitoring and motor planning during auditory-motor integration^30,58,64^. In CA, gray matter atrophy of IFG, impaired white matter integrity of the AF and unstable IFG-STG connectivity occurred in the right FTC jointly limit the circuit’s capacity to extract auditory information and relay motor plans, thereby constraining auditory-motor integration. Consequently, although the core deficit of CA may be improved by intervention^65,66^, such intervention remains inefficient. For conditions characterized by stable impairment with mechanistic constraints, practical efforts should therefore extend beyond remediating the impaired domain alone and instead strengthen the identification and cultivation of potential advantage domains.

### Potential competitive-like benefit in left FTC

**Structural benefits in left FTC.** Although no group differences were observed in the unilateral gray matter structure within the FTC, CA showed a competitive-like enhancement of leftward lateralization of IFG’s GMV (numerically smaller on the right and larger on the left than controls). Correlation analysis further revealed that the LI of IFG’s GMV was significantly negatively correlated with MBEA scores, supporting the hypothesis that the right FTC is impaired while the left achieves competitive gains rather than compensation. Specifically, reduced GMV of right IFG may create favorable conditions for the left homotopic region to acquire more neural resources during competition, ultimately leading to increased GMV of left IFG and a competitive-like enhancement of leftward lateralization. Regarding white matter microstructure, we found a significant leftward lateralization of FA in the dAF in CA. Unlike GMV, CA did not show a competitive-like alteration, with numerically lower FA in the left hemisphere. That is, the lateralization change was mainly driven by reduced FA in the right hemisphere. This phenomenon may be explained, on the one hand, interhemispheric interactions at the white matter microstructural level may be more complex, which may involve not only competitive changes but also adaptive up– or downregulation to maintain interhemispheric stability ^67,68^. On the other hand, microstructure of left dAF in CA may also have mild impairment; in this case, even if a competitive benefit arises, the inherent impairment may prevent the typical seesaw-like competitive pattern from being observed. Correlation analyses further showed that the enhanced leftward lateralization of FA in the dAF was negatively correlated with MBEA scores, which argues against a compensatory interpretation of this alteration. Instead, such changes may create favorable conditions for functional gain in the left FTC, which we discuss below.

**Functional communication benefits in left FTC.** For sFC, CA showed stronger connectivity between the left IFG and STG, and this enhancement was accompanied by a higher proportion of CA individuals with positive sFC deviations. Correlation analysis showed that stronger left IFG–STG sFC and its deviation score were negatively correlated with MBEA scores, indicating that the enhancement was not a compensatory response to the music deficits. Previous studies have shown that the functional activity level in one hemisphere depends not only on the integrity of its ipsilateral structural substrate, but also on the structural state of its contralateral homologue ^69–72^. For example, Vernooij et al. found that the integrity of the right AF was negatively correlated with the relative strength of functional activity in the left FTC, suggesting that reduced white matter integrity in one hemisphere may promote functional activity within the contralateral homologue by releasing competitive resources or weakening interhemispheric inhibition^73^. In the present study, impaired right AF and enhanced leftward lateralization of AF microstructure in CA may have weakened the inhibitory regulation of left-hemispheric functional activity. This may have released neural resources or created a “competitive vacuum” in the left homotopic circuit, thereby enhancing its functional connectivity. Therefore, the increased sFC of left FTC in CA more likely reflects a competitive benefit. Regarding dFC, similar to GMV, CA showed a competitive-like enhancement of leftward lateralization in dFC between the IFG and STG (numerically smaller on the left and larger on the right than in controls). In other words, CA showed enhanced leftward lateralization of functional connectivity stability between the IFG and STG. Stronger leftward lateralization of stability was negatively correlated with MBEA scores, suggesting that such changes may also reflect a competitive benefit arising from right FTC impairment rather than a compensatory response to the music deficits. Overall, these connectivity findings converge with the structural evidence to suggest that the left FTC in CA exhibits competitive benefits rather than compensation. One recent study reported reduced left IFG–STG connectivity in CA ^74^, whereas we observed the opposite pattern. This inconsistency is discussed in Supplementary Discussion 2.

**Heightened flexibility of the left IFG.** CA also showed stronger left IFG dALFF, which was negatively correlated with MBEA scores. Although dALFF and dFC are both dynamic metrics, they capture different levels of neural dynamics: dALFF measures the variability of spontaneous activity at a single node, whereas dFC measures fluctuations in inter-regional coupling at the network level. Their values therefore need not carry the same functional meaning. In the present study, greater dALFF is better interpreted as heightened flexibility, that is, a capacity to rapidly reconfigure local activity in response to changing demands. This fits the role of the left IFG in coordinating the transformation of information across large-scale language-related networks^75^, a function that inherently depends on flexibility. Consistent with this view, Guo et al. reported reduced dALFF of left IFG in aphasia, with greater reductions associated with more severe speech-production impairment ^76^. Moreover, focal nodal damage does not necessarily produce persistent impairment, whereas disrupted connectivity does^77^. Accordingly, even if elevated dALFF reflected less stable local activity, the relatively robust left FTC architecture, as evidenced by enhanced strength and stability of IFG-STG connection, may buffer this instability and preserve overall processing efficiency. Overall, the left FTC of CA may thus combine a highly flexible node with a stable and efficient network.

**Speech production advantages in CA.** Study 2 found that after strategy instruction, both groups produced more content (e.g., more TNW and TNU), showed greater lexical diversity, described more events accurately with fewer omissions and captured the story theme better, indicating that the narrative strategy was effective. However, these gains likely increased cognitive load, as participants had to monitor more events and details, organize content and maintain the narrative thread. This increased load may have pushed task demands beyond the available cognitive-neural resource capacity^78–80^, which in turn reduced fluency, shortened MLU and lowered local coherence and informativeness in both groups at post-test. Crucially, CA diverged from controls under this elevated load. CA maintained syntactic complexity across pre– and post-test, whereas controls showed a significant decline. CA also produced longer utterances and showed a higher proportion of individuals with developmental advantages in syntactic complexity and local coherence. In addition, CA spoke faster and showed higher informativeness efficiency while producing comparable narrative content. Together, these patterns indicate that CA can preserve or even enhance the speech quality under high cognitive load, pointing to advantages in speech planning, load coping and expressive efficiency. These characteristics may place CA favorably in open-ended speech tasks requiring real-time, rapid and accurate production of large amounts of factual information, such as live sports commentary and field reporter. These behavioral advantages also converge with the left FTC features of CA observed in Study 1. The anterior temporal regions support lexical and semantic processing and projects lexical representations to frontal regions^81,82^. Study 1 found enhanced functional connectivity stability between the left IFG and anterior temporal regions (e.g., STG.A38m and lateral area 38 of STG), which may support rapid lexical retrieval and stable input of lexical information into syntactic encoding. In addition, the dorsal frontotemporal pathway constituted by the left AF, connects Broca’s (IFG.A44v) and Wernicke’s areas (STG.A41/42, rpSTS and STG.A22c) and supports syntactic processing^81^. Study 1 found that CA showed increased strength and stability of functional coupling within the pathway, as well as stronger leftward lateralization of AF integrity. Together, these features may form a more efficient and robust syntactic-generation circuit, allowing CA to maintain syntactically more complex sentences under load. Efficient anterior temporal-frontal-posterior temporal connectivity may make syntactic processing more fluent and automatized, thereby conserving resources for higher-level narrative processing such as structuring the narrative and monitoring the narrative thread. These conserved resources may in turn support tighter semantic cohesion and smoother transitions, that is, better local coherence under complex conditions. Finally, heightened dALFF in the left IFG further supports this resource-based account and supports interpreting nodal dALFF as heightened flexibility, which enables rapid resource recruitment and redistribution for online speech planning and load coping under high cognitive load. Dissociation between objective advantages and subjective experience, and alternative explanations for such speech-production advantage are discussed in Supplementary Discussion 3.

### Pause pattern as a potential adaptive speech-planning strategy

Pauses in narrative speech serve multiple functions^83^. Beyond physiological needs such as breathing, they mark syntactic boundaries (e.g., short silent pauses typically separating clauses and medium pauses separating sentences^84^). Pauses also can serve narrative goals such as drawing listener attention, and help to buy time for the planning, execution, monitoring and repair of speech^85,86^. Long silent pauses commonly signal hesitation or blocked speech planning^84^, and in interactive settings they can create awkward silences^87^ or lead to loss of the floor through interruption^88^. In the present study, CA and controls produced comparable TNU, and since short and medium pauses served only syntactic marking here, their frequencies should in theory have been similar. Instead, CA showed more short silent pauses and poorer development of medium silent pauses, alongside fewer long silent pauses and more filled pauses. This pattern suggests that when CA needed more time for speech planning, they tended to use filled pauses rather than long silences. Filled pauses can secure additional planning time while maintaining the continuity of the speech stream, and they can signal to listeners that speech is ongoing, helping to avoid silences or interruption. By either replacing long silences with filled pauses or breaking them into shorter silent pauses, CA achieved a pattern of fewer long pauses but more filled and short silent pauses. This pattern may reflect an adaptive strategy suited to specific spoken interaction situations.

### Spared episodic memory as evidence for mechanism specificity

CA performed comparably to controls on both episodic memory tasks, this finding further indicates that the musical impairment of CA is mainly related to music memory deficits rather than a general memory impairment^89–91^. Episodic memory mainly depends on the medial temporal lobe/hippocampal network^92,93^, which is not the core impaired substrate in CA and is not predicted to benefit competitively. The absence of group differences therefore supports the specificity of the competitive-benefit mechanism, indicating that the advantage is confined to the circuit undergoing competitive reallocation instead of reflecting a general compensatory or global-enhancement account.

### Implications

Deficit-focused research generates advantages as by-products, to be discovered retrospectively and explained post hoc. The pathway tested here inverts that order. It specifies from cognitive neuroscience where an advantage should be sought, and then tests that prediction behaviorally. The prediction was falsifiable, and it held in the domains where an advantage was expected while correctly predicting its absence elsewhere. This makes the framework compatible with, rather than opposed to, conventional deficit assessment. It does not require that impairments be redefined as differences (like neurodiversity^4,94,95^), only that the neural basis of impairment be treated as a starting point for asking what else the system can do and not solely for asking what it cannot do. Its practical value is consequently straightforward, encompassing earlier and more targeted identification of strengths for educational and occupational guidance, support for letting individuals develop along the functions that remain relatively spared instead of directing their effort mainly toward constrained ones. And because strengths and impairments are shown to coexist within a unified cognitive neuroscience framework, the present study also provides an evidence-based basis for destigmatizing atypical populations.

### Limitations and Further Directions

The most important limitation is that the two studies were conducted in independent samples. Such design guards against sample-specific overfitting and mirrors the practical situation in which neuroimaging is unavailable, but we cannot link neuroimaging features with behavior directly. Although the two studies converge on the same prediction from different levels, studies combining multimodal MRI and behavior data in the same participants is the necessary next step. Second, although our findings align with competitive benefit, they do not license a causal claim that competition itself produced the advantages. Future research needs to verify the causal relationship by directly manipulating the competing circuits, or by tracking neural and behavioral development longitudinally from early life stage. Third, many variables could not be modelled with adequate normative performance, and the CA sample size in Study 2 was small. Future studies should identify more effective predictors for these measures so that normative models can be built more effectively, and should replicate to further verify current findings. Finally, whether this cognitive-neuroscience-based pathway for predicting cognitive advantages generalizes beyond CA to other mental conditions remains an open question and is the appropriate next test.

## Conclusion

The right FTC is impaired in CA, whereas the competing left FTC shows competitive-like neuroimaging benefits and supports an advantage in speech production, alongside spared episodic memory and constrained music learning. These findings provide empirical support for a cognitive-neuroscience-based pathway that can prospectively predict cognitive advantages, and suggest a practical route for translating basic cognitive-neuroscience findings into strength-based assessment and intervention.

## Methods

### Participants

Two independent samples were recruited from universities in Changsha, Hunan, China. Participants were included if they were right-handed and had no systematic musical training, and were excluded if they had hearing impairment, a history of brain injury, major neurological illness, or psychiatric illness, or a WAIS score below 85. Study 1 additionally excluded individuals with MRI contraindications, and Study 2 additionally excluded individuals with formal training in speech or narrative expression. CA was defined as an MBEA score ≤ 21.5. Finally, Study 1 recruited 69 CA and 97 controls (during November 2018 – December 2023). Study 2 recruited a separate sample of 30 CA and 103 controls (during December 2024–October 2025). Study 2 needed more controls because cognitive advantage was identified via normative modelling, which requires a large reference sample for stable estimation. Written informed consent was obtained from all participants, and both studies were approved by the Ethics Committee of the Second Xiangya Hospital, Central South University.

Regarding normative modelling for MRI, an independent reference sample was drawn from the INDI. After preliminary quality control, 1019, 1016, and 1002 subjects were retained for rs-fMRI, sMRI and dMRI respectively (1023 subjects in total). Dataset composition and basic information are reported in Supplementary Table 1; screening flow, hearing and IQ assessment, selection rules for INDI datasets were described in Supplementary Method 1.

Age distributions of Xiangya samples and INDI samples were presented in Supplementary Fig.1

### Background Measures

**MBEA.** MBEA^52^ comprises six subtests (scale, contour, interval, rhythm, meter and memory) of 30 valid items each. In the three pitch-related subtests (scale, contour and interval) and the rhythm subtest, participants judged whether each pair of musical fragments was the same or different. In the melody subtest, they judged whether the presented fragment was a waltz or a march. In the memory subtest, they judged whether a fragment had appeared in the preceding five subtests. One point was awarded per correct response, and the composite score was the mean of the six subtests. The composite score served both as the CA screening criterion (≤21.5^53^) and as the continuous index of music perception ability in brain–behavior analyses.

**WAIS.** We conducted the Chinese version of WAIS to estimate general intelligence. Participants scoring below 85 were excluded.

**Goldsmiths Musical Sophistication Index (Gold-MSI).** The Gold-MSI is a self-report questionnaire used to measure individual musical sophistication^96^. Its Chinese revised version comprises 38 items scored on a seven-point scale and yields five subscales (active engagement, perceptual abilities, music training, singing abilities, and emotions) as well as a general sophistication dimension^97^. Higher scores indicated better self-perceived ability. In the present study, the Cronbach’s α for this scale was 0.904.

**Self-perceived narrative expression.** This was measured with a self-developed eight-item questionnaire covering clarity, coherence, logic, confidence, appeal, others’ evaluations and willingness to speak in familiar and unfamiliar settings, scored on a five-point scale (Cronbach’s α = 0.789). Detailed item content and administration order are provided in Supplementary Method 2.

### ROI Definition of FTC

**IFG and STG.** The Brainnetome atlas^98^ was used to define the IFG and STG. The IFG ROI comprised labels 29–40, and the STG ROI comprised labels 69–80. We also considered labels 121–124 as part of the STG, corresponding to the posterior superior temporal sulcus in the Brainnetome atlas. Detailed information and anatomical locations for the above ROIs is provided in Supplementary Table. 2 and Supplementary Fig. 3.

**AF Segments and Seeds for Reconstruction.** The AF is the core white matter pathway within the FTC and is the tract most reported to be altered in CA. Following previous studies^15,37^, we divided the AF into three segments: anterior, posterior and direct. The anterior segment mainly connects the posterior frontal lobe (posterior IFG and ventral precentral gyrus) with the inferior parietal lobule (IPL), the posterior segment connects the IPL with the posterior temporal gyrus, and the direct segment connects the posterior frontal lobe (posterior IFG and ventral precentral gyrus) with the posterior temporal gyrus. Seed regions for AF reconstruction were defined with the Brainnetome atlas. Frontal seeds comprised labels 17/18, 29/30, 33/34, 39/40 and 63/64; temporal seeds comprised labels 71/72, 75/76, 121/122 and 123/124; parietal seeds comprised labels 137/138, 139/140 and 141/142. Detailed information for the ROIs comprising the seed regions is provided in Supplementary Table. 2. The resulting seed masks are shown in Supplementary Fig. 4a.

### Image acquisition and processing

Participants in Study 1 were scanned on a 3T Siemens Skyra scanner at the Second Xiangya Hospital, Central South University. Detailed acquisition parameters were presented in Supplementary Method 3. Imaging acquisition parameters of INDI are available from the dataset providers (See Supplementary Method 1).

rs-fMRI, sMRI and dMRI were preprocessed using DPABI, CAT12 and FSL respectively. For rs-fMRI, we calculated sALFF across all ROIs to reflect the intensity of spontaneous neural activity of core nodes within the FTC; we also calculated the sFC between each IFG ROI and its ipsilateral STG ROIs, as well as between all IFG and STG ROIs and their respective contralateral homologues, to reflect functional coordination within the FTC. To measure the variability of spontaneous neural activity intensity and functional coordination, we additionally calculated dALFF and dFC. Because dynamic metrics are sensitive to sliding-window parameters, we used a window length of 30 volumes as the primary analysis and 50 volumes as a validation analysis, with an 80% overlap between sliding windows in both cases. For sMRI, we calculated the GMV and cortical thickness of all ROIs to measure the grey matter morphology of core FTC nodes. For dMRI, we reconstructed sub-segments of the AF using probabilistic tractography, and then calculated the microstructural (including FA, MD, AD and RD) and macrostructural features (including volume and connection density) of each segment. Except the FC between IFG and its contralateral homologue, we also calculated the lateralization index (LI) for all metrics, expressed as (left – right) / (left + right), to measure the relative difference between homologous regions in the two hemispheres. Detailed image preprocessing and metric calculation procedures are provided in the Supplementary Method 3.

### TtL cognitive tasks design

In the behavioral assessment of cognitive advantage, an individual’s cognitive advantage can be defined by two complementary standards. Achievement refers to an individual’s performance level at a given time point, with above-norm performance indicating an achievement-level advantage. Development refers to the extent and upper limit of improvement following learning, training, or strategy use, with faster gains or a higher ceiling indicating a development-level advantage. Distinguishing between these two dimensions is necessary because relying solely on the achievement-based standard cannot accurately evaluate an individual’s cognitive advantage. Excellent cognitive performance at a single time point may merely stem from earlier cognitive training with a low growth ceiling, whereas poor performance may result from untapped potential or a lack of the necessary conditions to demonstrate talent. Therefore, we adopted TtL-adapted cognitive tasks, which allow the evaluation of cognitive advantages and potential across both achievement and development dimensions^99^.

**Speech production.** We conducted a storytelling task to elicit speech and assess speech production ability. The elicitation material was excerpted from *The Wanderings of Sanmao* (*San Mao Liu Lang Ji*, a classic Chinese comic), which comprised 30 panels with temporal and causal links forming seven episodes. The story concerns a wandering orphan who tries to change his circumstances, briefly finds companionship with an elderly man but is left alone again after the man is killed. Apart from panel numbers indicating sequence and years indicating the historical period, the material contained no textual cues about theme or plot. The task was administered twice to implement the TtL design, with standard narrative strategy instruction provided between the two administrations. In the pre-test, participants viewed the panels for 5 min to understand the story and organize their narration silently, and were then asked to tell the story as if to a child, as fully, vividly and fluently as possible, with no time limit. After the pre-test, they received standard narrative strategy guidance such as establishing a theme or storyline and using it to link events and sub-episodes. After guidance and another 5 min preparation, they narrated again under the same conditions (post-test). All the subjects’ narrations were digitally recorded and transcribed subsequently. Speech production performance were evaluated through fluency (including short/medium/long silent pauses frequency and duration, total silent pauses frequency and duration, filled pauses frequency, repetition frequency and repair frequency, speech rate, voiced pauses frequency and total pauses frequency), micro-linguistic (including TNW, TNU, lexical diversity, MLU and syntactic complexity) and macro-linguistic features (theme search, global and local coherence, event accuracy, informativeness density and efficiency). Detailed descriptions and calculation methods for the above measurements were provided in the Supplementary Method 5 and Supplementary Method 6.

**Episodic memory.** We assessed episodic memory using the AVLT^56^ and the VPRT^57^, both of which have been adapted to the TtL paradigm in previous studies. In AVLT, participants were presented with 15 two-character Chinese words across six recall trials. The first two trials served as the pre-test, trials 3 to 5 served as the practice phase with feedback and trial 6 served as the post-test. A delayed recall followed 10 min after trail 6. One point was awarded for each correctly recalled word, and the pre-test score was the mean of the first two trials. The VPRT comprised three blocks. In each block, participants first memorized 10 target pictures and then recognized them among 50 pictures. The 50 pictures comprised 10 targets, 20 highly similar lures and 20 dissimilar lures. No instruction was provided in the first block; after that, participants were given a mnemonic strategy (verbalizing out what is happening in each picture) and required to use it in the second block. In the third block, the strategy was not mandatory. All pictures were drawn from another classic Chinese comic named *Lao Fu Zi* (examples are provided in the Supplementary Method 7), and task performance was evaluated by the number of false recognitions.

**Rhythm Learning Task.** We assessed musical ability using a musical rhythm task adapted from previous studies^15,58^. The task comprised three blocks, each containing 12 trials. Each trial consisted of a listening phase and an imitation phase. During the listening phase, participants learned an 11-note rhythm sequence (lasted approximately 6 s) without using bodily movements to aid memorization. During the imitation phase, participants tapped the SPACE button of keyboard to synchronize with the rhythm as accurately as possible. A warning tone was presented 500 ms before each target sequence to signal the start of the trial. Participants were instructed not to stop and correct errors, but to continue following the rhythm even when uncertain. Before the formal learning began, participants completed two additional practice blocks with two rhythms not included in the learning protocol to familiarize themselves with task requirements. The rhythms used in the three blocks were composed of the same 11 notes, but different permutations of note order formed three distinct rhythms. Task performance was evaluated by accuracy and MAI. For each block, the first three trials served as the pre-test and the last three trials served as the post-test. Detailed information on rhythm composition and indicator calculation is provided in the Supplementary Method 7.

### Normative modelling and individual deviation

Normative modeling is a statistical technique that builds a reference model of how response variables of interest (e.g., GMV) vary with clinically relevant variables (e.g., age and sex), much like pediatric growth charts that plot a child’s height or weight against age^100,101^. It then allows each patient to be placed against this reference to quantify how far and in what direction they deviate from the expected normative range, thereby enabling assessment at the individual level rather than only at the level of group means and thus capturing heterogeneous, individual-specific patterns that group averages tend to obscure.

In Study 1, we used Hierarchical Bayesian Regression (HBR) in PCNtoolkit to fit normative model, because HBR could deal with site effect while avoiding the over-regression caused by harmonization methods such as ComBat^102^. Age, sex, dataset label and modality-specific covariates (mean FD for rs-fMRI; total intracranial volume and IQR for sMRI) were used as predictors for each neuroimaging feature to construct normative models and calculate deviation scores. The calculation formula for the deviation score z is:

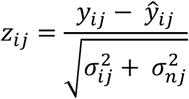

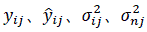 denote the actual measured value and predicted value of the neuroimaging feature of the j-th brain region for individual i, the variance of the measured value of the neuroimaging feature of the j-th brain region for individual i (prediction uncertainty) and the prediction variability of the model on the measured value of the neuroimaging feature of the j-th brain region, respectively.

INDI participant data were stratified by source dataset and randomly split 8:2 into training and test sets. Basic models were built on the training set and its generalization was evaluated on the test set. Basic models with explained variance > 0 were then transferred by stratifying CA and control participants by group label and randomly dividing them 1:1 into two subsets, one used to adapt the baseline model to the new-site data and the other used to derive individual deviation scores. The two subsets were then swapped and the process repeated to generate deviation scores for the participants in the original training subset, and the results from both rounds were combined to produce the final complete set of deviation scores for all CA and control participants. For each neuroimaging feature, individuals were categorized as showing positive or negative deviation based on cutoff values of the deviation score z = 0 (deviation direction) or |z| = 1.96 (extreme deviation).

In Study 2, we conducted Gaussian Process Regression (GPR) in PCNtoolkit to fit the normative models. For the achievement level, models were built using controls’ pre-test data with sex, age, education years, and WAIS score as predictors, and leave-one-out cross-validation (LOOCV) was used to evaluate the models and generate deviation scores for controls’ pre-test data; models with explained variance > 0 were then applied to controls’ post-test data and each stage’s data of the CA group to compute deviation scores. Similarly, for developmental-level normative modelling, pre-test scores were additionally included as predictors to build normative models for post-test performance from controls’ data, and LOOCV was used to evaluate the models and generate deviation scores; models with explained variance > 0 were then applied to the CA group to compute deviation scores, which quantify training gains relative to each individual’s baseline and demographic profile, thereby identifying potential cognitive advantage on the developmental dimension. We didn’t build normative models for global coherence and theme search because the range of both measure is limited. Individuals were considered to show advantages if they had a positive deviation score on measures where higher scores represent better performance, or a negative deviation score on measures where lower scores represent better performance; otherwise, they were considered to show disadvantages. The selection of deviation score thresholds was the same as in Study 1.

### Statistical analysis

Group differences in demographic, non-TtL cognitive metrics, neuroimaging metrics and the proportions of individuals classified as positive/negative deviation were tested with independent-samples t tests or χ² tests. Correlation between MBEA scores and neuroimaging features showing group differences were assessed with partial correlations controlling sex, age, education years, mean FD, IQR and total intracranial volume where applicable; Pearson correlations were used for correlation between MBEA score and deviation scores. Bonferroni correction was applied for multiple comparison correction of group comparison of neuroimaging features. TtL task performance was analyzed using (generalized) MLM, with Group, Time and their interaction as fixed effects, and participant as a random effect; sex, age, education or WAIS score were included as covariates when they differed between groups and were associated with the dependent variable. Simple-effects analyses were used to decompose significant interactions with Bonferroni correction.

## Data availability

INDI data are available from the website https://fcon_1000.projects.nitrc.org/indi/IndiRetro.html and https://fcon_1000.projects.nitrc.org/indi/IndiPro.html. The Xiangya data (including neuroimages in Study 1 and behavioral data in Study 2) are not available for public download, but access can be requested through the corresponding author D.W. Requests for raw and analyzed data will be promptly reviewed by the ethics committee of the Second Xiangya Hospital to verify whether the request is subject to any intellectual property or confidentiality obligations.

## Code availability

The PCN toolkit (https://pcntoolkit.readthedocs.io/en/v0.35/index.html) was used to build normative models and generate deviation scores. Transcription of recordings was performed using the large-model automatic speech recognition API provided by Volcengine (https://www.volcengine.com/docs/6561/1354868). Natural language analysis was conducted with HanLP (https://github.com/hankcs/HanLP), LTP (https://github.com/HIT-SCIR/ltp), the reasoning mode of DeepSeek-V3.2 (https://api-docs.deepseek.com/news/news251201) and text-embedding-v4 (https://help.aliyun.com/zh/model-studio/embedding).

## Acknowledgements

This work was partly supported by a research grant from the National Natural Science Foundation of China awarded to D.W. (No. 81771172). We acknowledge that the INDI datasets used in the present study were supported by multiple funding sources: the Beijing Enhanced and Beijing Short TR datasets were supported by a grant from the National Natural Science Foundation of China (No. 30770594) and a grant from the National High Technology Program of China (863 Program; No. 2008AA02Z405); the SLIM dataset was supported by the National Natural Science Foundation of China (Nos. 31271087, 31470981, 31571137 and 31500885), the National Outstanding Young People Plan Program for the Top Young Talents by Chongqing, the Fundamental Research Funds for the Central Universities (SWU1509383 and SWU1509451), the Natural Science Foundation of Chongqing (cstc2015jcyjA10106), the Fok Ying Tung Education Foundation (151023), the General Financial Grant from the China Postdoctoral Science Foundation (2015M572423 and 2015M580767), the Special Funds from the Chongqing Postdoctoral Science Foundation (Xm2015037), and the Key Research for Humanities and Social Sciences of Ministry of Education (14JJD880009). The National Institute on Drug Abuse and the National Natural Science Foundation of China (NSFC) have been instrumental in the CoRR collaboration, providing the necessary funding and manpower to build the foundation of the project, along with the Child Mind Institute, the Institute of Psychology, Chinese Academy of Sciences, and the Nathan Kline Institute. We thank all individuals and institutions that have contributed to the construction of the aforementioned databases. We would like to thank Bowen Ran, Dongling Yuan, Bochen Wang, Chenhan Xu, Yu Song, Yusheng Huang and Fanchen Meng for their valuable assistance during the experimental design and preparation phase.

## Author contributions

Y.W. conceived the study. Y.W. and M.L. wrote the manuscript. Y.W. performed the formal analysis and interpreted the results, with contributions from M.L. The following contributed to conduct the experiment: Y.W., Z.J., S.Huyang, N.G., J.Z., Q.L., C.S., Q.L., S.Huang and J.Y. D.W. secured funding, provided resources and administered the study. All authors reviewed the manuscript.

## Competing interests

All authors declare no competing interests.

**Extended Data Fig. 1.**
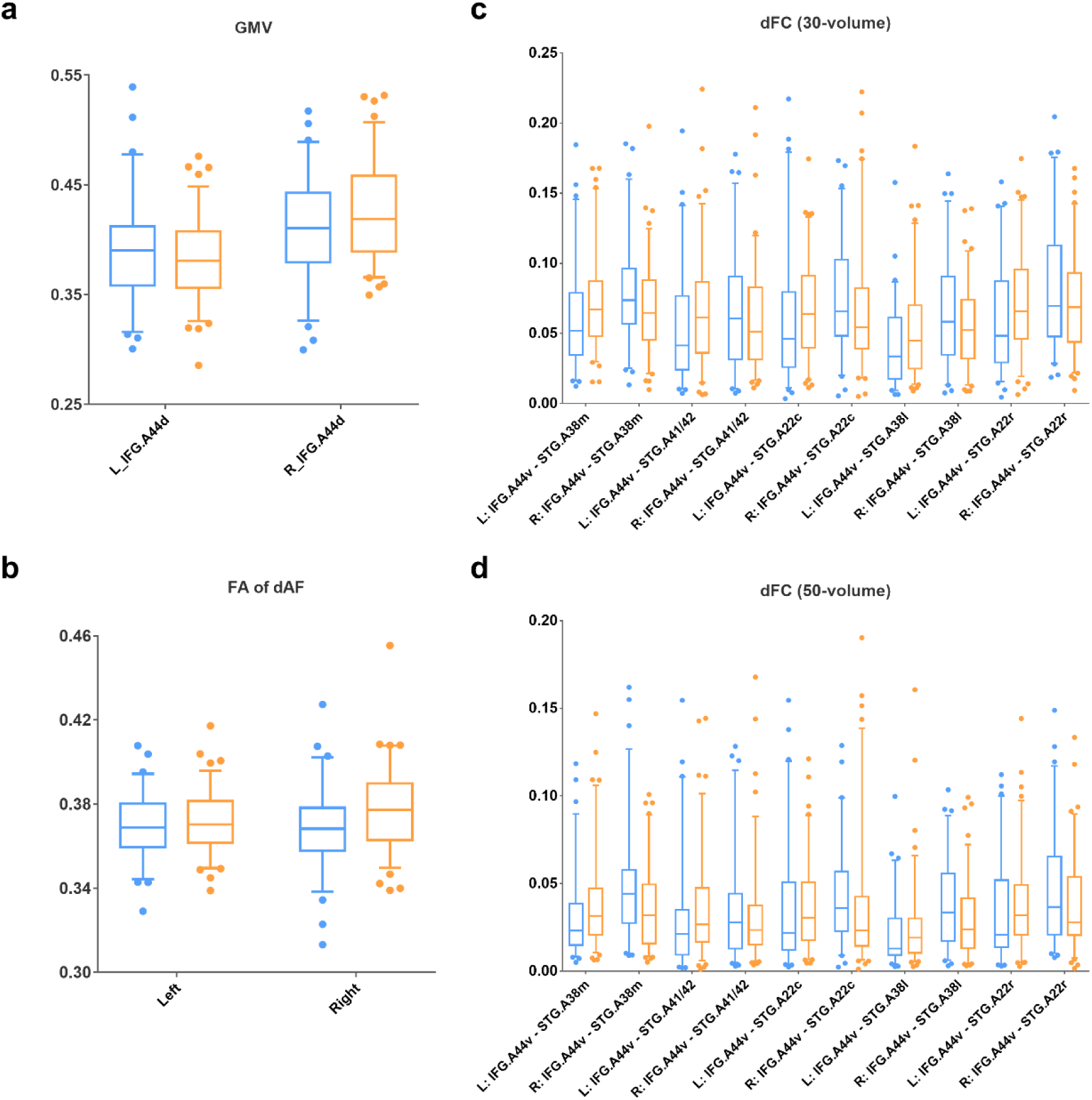
Box-plots of raw values for each hemisphere in MRI metrics showing significant group differences in lateralization index. **a**, congenital amusia showed numerically higher GMV in the left and numerically lower GMV in the right IFG.A44d. b, congenital amusia showed numerically lower FA in both left and right dAF. **c-d,** congenital amusia showed numerically higher dFC in the right and numerically lower dFC in the left. Orange represents controls and blue represents congenital amusia. The upper and lower bars represent 95 and 5 percentile, respectively. GMV, grey matter volume; sFC, static functional connectivity; FA, fractional anisotropy; dFC, dynamic functional connectivity; IFG, inferior frontal gyrus; STG, superior temporal gyrus; A44d, dorsal area 44; A41/42, area 41/42; A44v, ventral area 44; A45c, caudal area 45; A38m, medial area 38; A22c, caudal area 22; A38l, lateral area 38; A22r, rostral area 22; dAF, direct arcuate fasciculus; R, right; L, left

**Extended Data Table. 1.** Demographic comparisons between both groups in Study 1 (M ± SD)

|  | CA | Controls | χ<sup>2</sup>/t | *p* | Cohen’s d |
|---|---|---|---|---|---|
| Sex (Male/Female) | 32/37 | 32/65 | 3.050 | 0.081 | - |
| Age (Years) | 19.51 ± 1.90 | 19.93 ± 1.89 | -1.408 | 0.161 | -0.222 |
| Education Years | 13.65 ± 1.20 | 14.11 ± 1.66 | -1.967 | 0.051 | -0.310 |
| WAIS | 108.48 ± 9.36 | 109.20 ± 9.02 | -0.497 | 0.620 | -0.078 |
| MBEA |  |  |  |  |  |
| Scale | 19.20 ± 2.74 | 26.52 ± 2.95 | -16.224 | < 0.001 | -2.555 |
| Contour | 19.24 ± 3.39 | 27.37 ± 2.20 | -18.721 | < 0.001 | -2.948 |
| Interval | 18.22 ± 3.49 | 25.96 ± 3.31 | -14.528 | < 0.001 | -2.288 |
| Rhythm | 19.90 ± 3.81 | 26.44 ± 3.09 | -12.202 | < 0.001 | -1.922 |
| Meter | 18.15 ± 4.03 | 23.81 ± 4.02 | -8.944 | < 0.001 | -1.409 |
| Memory | 20.10 ± 4.22 | 27.69 ± 2.14 | -15.197 | < 0.001 | -2.393 |
| Global Mean | 19.14 ± 1.84 | 26.30 ± 2.01 | -23.441 | < 0.001 | -3.692 |
| Mean FD | 0.07 ± 0.03 | 0.07 ± 0.03 | -0.538 | 0.591 | -0.087 |
| TIV | 1505.93 ± 148.57 | 1487.02 ± 123.35 | 0.872 | 0.384 | 0.140 |
| IQR | 2.10 ± 0.08 | 2.11 ± 0.08 | -0.532 | 0.596 | -0.086 |

Note: All non-MRI data in the table were calculated based on all Xiangya participants; all MRI-related variables were calculated based on the participants included in the formal analysis. Within the subset of participants included in the sMRI analysis, age (CA: 19.34 ± 1.51, controls 19.96 ± 1.92; *p* = 0.032) and years of education (CA: 13.61 ± 1.18, controls: 14.12 ± 1.65; *p* = 0.025) differed significantly between groups. CA, congenital amusia; TIV, total intracranial volume; M, mean; SD, standard deviation; WAIS, Wechsler Adult Intelligence Scale; MBEA, Montreal Battery of Evaluation of Amusia; FD, frame-wise displacement; IQR, image quality rating.

**Extended Data Table. 2.** Group-mean comparisons of MRI metrics (M ± SD)

| Metrics | CA | Controls | t | *p* | Cohen’s d |
|---|---|---|---|---|---|
| sFC |  |  |  |  |  |
| L_IFG.A44v – L_STG.A41/42 | 0.59 ± 0.24 | 0.48 ± 0.23 | 2.988 | 0.003 | 0.482 |
| L_IFG.A44v – L_rpSTS | 0.52 ± 0.28 | 0.39 ± 0.25 | 3.063 | 0.003 | 0.494 |
| dALFF |  |  |  |  |  |
| L_IFG.A45c (30-volume) | 0.32 ± 0.09 | 0.28 ± 0.07 | 3.199 | 0.002 | 0.516 |
| L_IFG.A44v (30-volume) | 0.35 ± 0.08 | 0.30 ± 0.06 | 4.085 | < 0.001 | 0.659 |
| L_IFG.A44v (50-volume) | 0.25 ± 0.06 | 0.21 ± 0.05 | 3.845 | < 0.001 | 0.620 |
| dFC-LI |  |  |  |  |  |
| IFG.A44v – STG.A38m (50-volume) | -0.21 ± 0.31 | 0.04 ± 0.36 | -4.619 | < 0.001 | -0.745 |
| IFG.A44v – STG.A41/42 (50-volume) | -0.12 ± 0.42 | 0.07 ± 0.37 | -3.067 | 0.003 | -0.495 |
| IFG.A44v - STG.A22c (50-volume) | -0.15 ± 0.42 | 0.04 ± 0.40 | -2.884 | 0.005 | -0.465 |
| IFG.A44v - STG.A38l (50-volume) | -0.25 ± 0.38 | -0.06 ± 0.39 | -2.924 | 0.004 | -0.472 |
| IFG.A44v - STG.A22r (50-volume) | -0.19 ± 0.39 | 0.02 ± 0.36 | -3.437 | < 0.001 | -0.555 |
| IFG.A44v - STG.A38m (30-volume) | -0.16 ± 0.29 | 0.05 ± 0.27 | -4.653 | < 0.001 | -0.751 |
| IFG.A44v - STG.A41/42 (30-volume) | -0.11 ± 0.36 | 0.05 ± 0.33 | -2.941 | 0.004 | -0.475 |
| IFG.A44v - STG.A22c (30-volume) | -0.14 ± 0.34 | 0.01 ± 0.31 | -2.952 | 0.004 | -0.476 |
| IFG.A44v - STG.A38l (30-volume) | -0.24 ± 0.31 | -0.05 ± 0.32 | -3.704 | < 0.001 | -0.598 |
| IFG.A44v - STG.A22r (30-volume)<sup>a</sup> | -0.20 ± 0.32 | -0.00 ± 0.29 | -3.948 | < 0.001 | -0.637 |
| GMV-LI of IFG.A44d | -0.02 ± 0.06 | -0.05 ± 0.06 | 3.088 | 0.002 | 0.497 |
| FA of R.dAF | 0.37 ± 0.02 | 0.38 ± 0.02 | -2.713 | 0.007 | -0.430 |
| FA of R.aAF | 0.35 ± 0.01 | 0.36 ± 0.01 | -2.730 | 0.005 | -0.432 |
| FA-LI of dAF<sup>b</sup> | 0.00 ± 0.02 | -0.01 ± 0.02 | 2.426 | 0.016 | 0.384 |

Note: Superscript a, the mean LI of dFC in the controls was specifically -0.0027. Superscript b, the mean LI of FA in the dAF in CA was specifically 0.0011. CA, congenital amusia; LI, lateralization index; GMV, grey matter volume; sFC, static functional connectivity; dALFF, dynamic amplitude of low-frequency fluctuations; FA, fractional anisotropy; dFC, dynamic functional connectivity; IFG, inferior frontal gyrus; STG, superior temporal gyrus; A44d, dorsal area 44; A41/42, area 41/42; A44v, ventral area 44; A45c, caudal area 45; A38m, medial area 38; A22c, caudal area 22; A38l, lateral area 38; A22r, rostral area 22; rpSTS, rostral posterior superior temporal sulcus; aAF, anterior arcuate fasciculus; dAF, direct arcuate fasciculus; R, right; L, left; M, mean; SD, standard deviation.

**Extended Data Table. 3.**
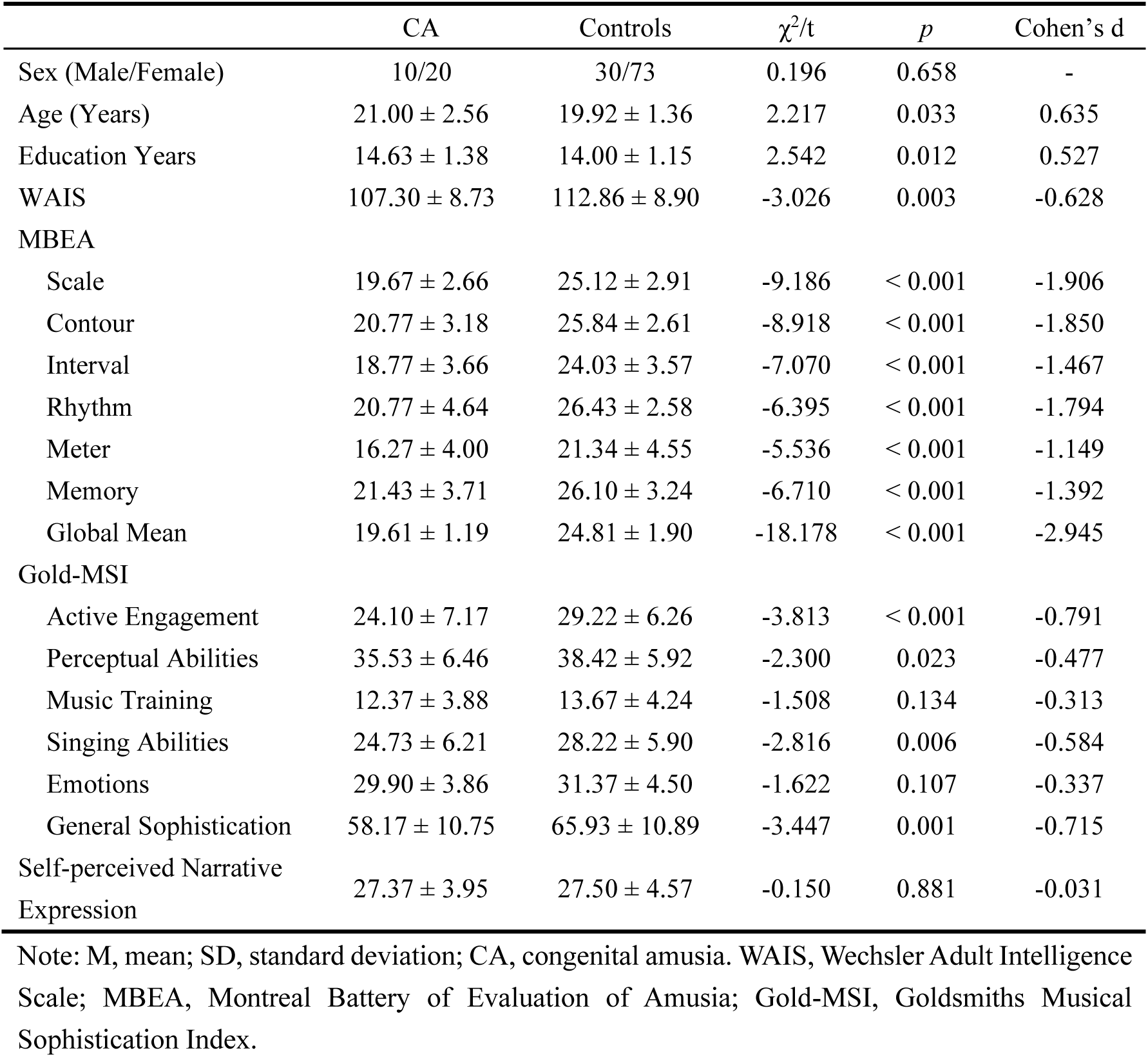
Comparisons of basic characteristics between both groups in Study 2 (M ± SD)

|  | CA | Controls | χ<sup>2</sup>/t | *p* | Cohen’s d |
|---|---|---|---|---|---|
| Sex (Male/Female) | 10/20 | 30/73 | 0.196 | 0.658 | - |
| Age (Years) | 21.00 ± 2.56 | 19.92 ± 1.36 | 2.217 | 0.033 | 0.635 |
| Education Years | 14.63 ± 1.38 | 14.00 ± 1.15 | 2.542 | 0.012 | 0.527 |
| WAIS | 107.30 ± 8.73 | 112.86 ± 8.90 | -3.026 | 0.003 | -0.628 |
| MBEA |  |  |  |  |  |
| Scale | 19.67 ± 2.66 | 25.12 ± 2.91 | -9.186 | < 0.001 | -1.906 |
| Contour | 20.77 ± 3.18 | 25.84 ± 2.61 | -8.918 | < 0.001 | -1.850 |
| Interval | 18.77 ± 3.66 | 24.03 ± 3.57 | -7.070 | < 0.001 | -1.467 |
| Rhythm | 20.77 ± 4.64 | 26.43 ± 2.58 | -6.395 | < 0.001 | -1.794 |
| Meter | 16.27 ± 4.00 | 21.34 ± 4.55 | -5.536 | < 0.001 | -1.149 |
| Memory | 21.43 ± 3.71 | 26.10 ± 3.24 | -6.710 | < 0.001 | -1.392 |
| Global Mean | 19.61 ± 1.19 | 24.81 ± 1.90 | -18.178 | < 0.001 | -2.945 |
| Gold-MSI |  |  |  |  |  |
| Active Engagement | 24.10 ± 7.17 | 29.22 ± 6.26 | -3.813 | < 0.001 | -0.791 |
| Perceptual Abilities | 35.53 ± 6.46 | 38.42 ± 5.92 | -2.300 | 0.023 | -0.477 |
| Music Training | 12.37 ± 3.88 | 13.67 ± 4.24 | -1.508 | 0.134 | -0.313 |
| Singing Abilities | 24.73 ± 6.21 | 28.22 ± 5.90 | -2.816 | 0.006 | -0.584 |
| Emotions | 29.90 ± 3.86 | 31.37 ± 4.50 | -1.622 | 0.107 | -0.337 |
| General Sophistication | 58.17 ± 10.75 | 65.93 ± 10.89 | -3.447 | 0.001 | -0.715 |
| Self-perceived Narrative Expression | 27.37 ± 3.95 | 27.50 ± 4.57 | -0.150 | 0.881 | -0.031 |

Note: M, mean; SD, standard deviation; CA, congenital amusia. WAIS, Wechsler Adult Intelligence Scale; MBEA, Montreal Battery of Evaluation of Amusia; Gold-MSI, Goldsmiths Musical Sophistication Index.

**Extended Data Table. 4.** Fluency performance (M ± SD) in the speech production task and linear mixed model comparisons.

| Metric | CA | Controls | Statistics |
|---|---|---|---|
| **Speech Rate** |  |  |  |
| Pre-test | 3.34 ± 0.64 | 3.01 ± 0.64 | Group：*F*<sub>(1, 131.000)</sub> = 6.848<sup>\*</sup><br>Time：*F*<sub>(1, 131.000)</sub> = 0.312<br>Group × Time：*F*<sub>(1, 131.000)</sub> = 0.026 |
| Post-test | 3.32 ± 0.51 | 3.00 ± 0.62 |  |
| **Frequency of Short Silence Pause** |  |  |  |
| Pre-test | 19.83 ± 6.68 | 17.81 ± 4.85 | Group：*F*<sub>(1, 131.000)</sub> = 4.060<sup>\*</sup><br>Time：*F*<sub>(1, 131.000)</sub> = 3.144<br>Group × Time：*F*<sub>(1, 131.000)</sub> = 0.301×10<sup>-3</sup> |
| Post-test | 20.49 ± 5.22 | 18.46 ± 4.94 |  |
| **Proportion of Short Silence Paused Duration** |  |  |  |
| Pre-test | 7.93 ± 2.72 | 7.01 ± 1.88 | Group：*F*<sub>(1, 131.000)</sub> = 4.943<sup>\*</sup><br>Time：*F*<sub>(1, 131.000)</sub> = 0.942<br>Group × Time：*F*<sub>(1, 131.000)</sub> = 0.152 |
| Post-test | 8.02 ± 2.07 | 7.21 ± 1.88 |  |
| **Frequency of Medium Silence Pause** |  |  |  |
| Pre-test | 9.58 ± 2.42 | 9.10 ± 2.53 | Group：*F*<sub>(1, 131.000)</sub> = 3.002<br>Time：*F*<sub>(1, 131.000)</sub> = 0.916<br>Group × Time：*F*<sub>(1, 131.000)</sub> = 1.112 |
| Post-test | 10.08 ± 2.44 | 9.07 ± 2.24 |  |
| **Proportion of Medium Silence Paused Duration** |  |  |  |
| Pre-test | 10.08 ± 2.55 | 9.77 ± 2.87 | Group：*F*<sub>(1, 131.000)</sub> = 1.354<br>Time：*F*<sub>(1, 131.000)</sub> = 0.541<br>Group × Time：*F*<sub>(1, 131.000)</sub> = 0.640 |
| Post-test | 10.51 ± 2.45 | 9.75 ± 2.41 |  |
| **Frequency of Long Silence Pause** |  |  |  |
| Pre-test | 3.83 ± 2.31 | 4.84 ± 2.49 | Group：*F*<sub>(1, 131.000)</sub> = 5.009<sup>\*</sup><br>Time：*F*<sub>(1, 131.000)</sub> = 0.545<br>Group × Time：*F*<sub>(1, 131.000)</sub> = 0.387 |
| Post-test | 3.85 ± 2.06 | 5.06 ± 2.66 |  |
| **Proportion of Short Silence Paused Duration** |  |  |  |
| Pre-test | 11.50 ± 8.91 | 15.79 ± 11.27 | Group：*F*<sub>(1, 131.000)</sub> = 5.015<sup>\*</sup><br>Time：*F*<sub>(1, 130.983)</sub> = 0.006<br>Group × Time：*F*<sub>(1, 130.983)</sub> = 0.507 |
| Post-test | 11.11 ± 6.82 | 16.27 ± 11.21 |  |
| **Frequency of Total Silence Pause** |  |  |  |
| Pre-test | 33.25 ± 7.12 | 31.75 ± 4.94 | Group：*F*<sub>(1, 131.000)</sub> = 2.657<br>Time：*F*<sub>(1, 131.000)</sub> = 6.870<sup>\*\*</sup><br>Group × Time：*F*<sub>(1, 131.000)</sub> = 0.196 |
| Post-test | 34.43 ± 5.30 | 32.59 ± 4.95 |  |
| **Proportion of Short Silence Paused Duration** |  |  |  |
| Pre-test | 29.51 ± 8.26 | 32.57 ± 9.55 | Group：*F*<sub>(1, 131.000)</sub> = 3.267<br>Time：*F*<sub>(1, 131.000)</sub> = 0.719<br>Group × Time：*F*<sub>(1, 131.000)</sub> = 0.326 |
| Post-test | 29.64 ± 6.79 | 33.24 ± 9.55 |  |
| **Frequency of Filled Pause** |  |  |  |
| Pre-test | 2.92 ± 3.20 | 1.41 ± 1.87 | Group：*F*<sub>(1, 129.373)</sub> = 9.296<sup>\*\*</sup><br>Time：*F*<sub>(1, 131.000)</sub> = 0.365<br>Group × Time：*F*<sub>(1, 131.000)</sub> = 0.025 |
| Post-test | 2.86 ± 3.47 | 1.31 ± 2.00 |  |
| **Frequency of non-Filled Pause** |  |  |  |
| Pre-test | 3.24 ± 3.57 | 2.41 ± 1.59 | Group：*F*<sub>(1, 129.037)</sub> = 1.187<br>Time：*F*<sub>(1, 130.990)</sub> = 0.363<br>Group × Time：*F*<sub>(1, 130.990)</sub> = 0.120 |
| Post-test | 3.37 ± 3.60 | 2.44 ± 1.73 |  |
| **Frequency of Voiced Pause** |  |  |  |
| Pre-test | 6.15 ± 4.92 | 3.82 ± 2.76 | Group：*F*<sub>(1, 129.338)</sub> = 7.730<sup>\*\*</sup><br>Time：*F*<sub>(1, 131.000)</sub> = 0.147×10<sup>-3</sup><br>Group × Time：*F*<sub>(1 ,131.000)</sub> = 0.102 |
| Post-test | 6.23 ± 5.28 | 3.76 ± 2.98 |  |
| **Frequency of Total Pause** |  |  |  |
| Pre-test | 39.40 ± 10.29 | 35.57 ± 6.31 | Group：*F*<sub>(1, 130.998)</sub> = 7.914<sup>\*\*</sup><br>Time：*F*<sub>(1, 131.000)</sub> = 4.522<sup>\*</sup><br>Group × Time：*F*<sub>(1 ,131.000)</sub> = 0.253 |
| Post-test | 40.65 ± 9.45 | 36.34 ± 6.57 |  |

Note: M, mean; SD, standard deviation. CA, congenital amusia. Speech rate is expressed in syllables per second. Pause frequency is expressed in number of pauses per minute. Proportion is presented as percentages.
\* *p* < 0.05; \*\* *p* < 0.01

**Extended Data Table. 5.** Micro-linguistic performance (M ± SD) in the speech production task and linear mixed model comparisons.

| Metric | CA | Controls | Statistics |
|---|---|---|---|
| **Total Number of Words** |  |  |  |
| Pre-test | 400.13 ± 120.16 | 428.65 ± 150.22 | Group： *F*<sub>(1, 130.000)</sub> = 0.001<br>Time： *F*<sub>(1, 131.000)</sub> = 163.935<sup>\*\*\*</sup><br>Group × Time： *F*<sub>(1, 131.000)</sub> = 0.777 |
| Post-test | 580.00 ± 210.08 | 585.35 ± 234.11 |  |
| **Total Number of Utterances** |  |  |  |
| Pre-test | 20.40 ± 7.27 | 24.06 ± 9.26 | Group： *F*<sub>(1, 130.000)</sub> = 1.112<br>Time： *F*<sub>(1, 131.000)</sub> = 117.825<sup>\*\*\*</sup><br>Group × Time： *F*<sub>(1, 131.000)</sub> = 0.008 |
| Post-test | 31.60 ± 13.16 | 35.08 ± 15.22 |  |
| **Lexical Diversity** |  |  |  |
| Pre-test | 60.20 ± 4.68 | 62.14 ± 3.86 | Group： *F*<sub>(1, 130.000)</sub> = 2.725<br>Time： *F*<sub>(1, 131.000)</sub> = 54.731<sup>\*\*\*</sup><br>Group × Time： *F*<sub>(1, 131.000)</sub> = 0.655 |
| Post-test | 62.53 ± 2.95 | 64.01 ± 3.42 |  |
| **Mean Length of Utterance** |  |  |  |
| Pre-test | 20.53 ± 5.34 | 18.45 ± 4.30 | Group： *F*<sub>(1, 131)</sub> = 4.831<sup>\*</sup><br>Time： *F*<sub>(1, 131.000)</sub> = 8.254<sup>\*\*</sup><br>Group × Time： *F*<sub>(1, 131)</sub> = 0.434 |
| Post-test | 18.99 ± 3.52 | 17.48 ± 4.58 |  |
| **Syntactic Complexity** |  |  |  |
| Pre-test | 52.56 ± 14.00 | 54.08 ± 12.30 | Group： *F*<sub>(1, 131.000)</sub> = 0.511<br>Time： *F*<sub>(1, 131.000)</sub> = 6.271<sup>\*</sup><br>Group × Time： *F*<sub>(1, 131.000)</sub> = 4.602<sup>\*</sup> |
| Post-test | 52.05 ± 10.00 | 47.47 ± 12.55 |  |

Note: M, mean; SD, standard deviation. CA, congenital amusia. Lexical diversity is computed using moving average type-token ratio method, and is expressed as percentages. Mean length of utterance is expressed in words per utterance. Syntactic complexity is presented as percentages. Total number of words and total number of utterances are expressed in counts.
\* *p* < 0.05; \*\* *p* < 0.01; \*\*\* *p* < 0.001

**Extended Data Table. 6.** Macro-linguistic performance (M ± SD) in the speech production task and linear mixed model comparisons.

| Metric | CA | Controls | Statistics |
|---|---|---|---|
| **Global Coherence** |  |  |  |
| Pre-test | 5/25 | 19/84 | Group：*F*<sub>(1, 262)</sub> = 0.271<br>Time：*F*<sub>(1, 262)</sub> = 23.944<sup>\*\*\*</sup><br>Group × Time：*F*<sub>(1, 262)</sub> = 0.054 |
| Post-test | 14/16 | 55/48 |  |
| **Theme Search** |  |  |  |
| Pre-test | 2.37 ± 1.00 | 2.54 ± 0.86 | Group：*F*<sub>(1, 131.000)</sub> = 0.849<br>Time：*F*<sub>(1, 131.000)</sub> = 35.993<sup>\*\*\*</sup><br>Group × Time：*F*<sub>(1, 131.000)</sub> = 0.070 |
| Post-test | 3.07 ± 1.01 | 3.18 ± 0.98 |  |
| **Local Coherence** |  |  |  |
| Pre-test | 0.53 ± 0.04 | 0.53 ± 0.05 | Group：*F*<sub>(1, 131.000)</sub> = 0.821<br>Time：*F*<sub>(1, 131.000)</sub> = 11.183<sup>\*\*</sup><br>Group × Time：*F*<sub>(1, 131.000)</sub> = 1.949 |
| Post-test | 0.52 ± 0.04 | 0.50 ± 0.05 |  |
| **Informativeness Density** |  |  |  |
| Pre-test | 50.44 ± 4.20 | 51.56 ± 4.49 | Group：*F*<sub>(1, 131.000)</sub> = 0.591<br>Time：*F*<sub>(1, 131.000)</sub> = 24.299<sup>\*\*\*</sup><br>Group × Time：*F*<sub>(1, 131.000)</sub> = 1.582 |
| Post-test | 48.92 ± 3.87 | 49.01 ± 4.24 |  |
| **Informativeness Efficiency** |  |  |  |
| Pre-test | 70.21 ± 14.16 | 64.83 ± 14.64 | Group：*F*<sub>(1, 131.000)</sub> = 4.981<sup>\*</sup><br>Time：*F*<sub>(1, 131.000)</sub> = 15.418<sup>\*\*\*</sup><br>Group × Time：*F*<sub>(1, 131.000)</sub> = 0.449 |
| Post-test | 67.19 ± 10.76 | 60.58 ± 13.29 |  |
| **Narrative Content (Accurate)** |  |  |  |
| Pre-test | 3.93 ± 2.36 | 5.23 ± 2.79 | Group：*F*<sub>(1, 128.000)</sub> = 0.160<br>Time：*F*<sub>(1, 131.000)</sub> = 33.210<sup>\*\*\*</sup><br>Group × Time：*F*<sub>(1, 131.000)</sub> = 1.042 |
| Post-test | 5.60 ± 3.06 | 6.40 ± 3.30 |  |
| **Narrative Content (Partially Accurate)** |  |  |  |
| Pre-test | 12.53 ± 2.57 | 12.13 ± 2.42 | Group：*F*<sub>(1, 131.000)</sub> = 0.389<br>Time：*F*<sub>(1, 131.000)</sub> = 2.886<br>Group × Time：*F*<sub>(1, 131.000)</sub> = 0.222 |
| Post-test | 11.97 ± 3.18 | 11.81 ± 2.42 |  |
| **Narrative Content (Miss)** |  |  |  |
| Pre-test | 8.47 ± 3.18 | 7.66 ± 2.86 | Group：*F*<sub>(1, 131.000)</sub> = 1.034<br>Time：*F*<sub>(1, 131.000)</sub> = 7.938<sup>\*\*</sup><br>Group × Time：*F*<sub>(1, 131.000)</sub> = 0.588 |
| Post-test | 7.53 ±3.32 | 7.13 ± 3.33 |  |
| **Narrative Content (Error)** |  |  |  |
| Pre-test | 5.07 ± 1.62 | 4.98 ± 2.20 | Group：*F*<sub>(1, 130.000)</sub> = 0.551<br>Time：*F*<sub>(1, 131.000)</sub> = 1.318<br>Group × Time：*F*<sub>(1, 131.000)</sub> = 0.120 |
| Post-test | 4.90 ± 2.17 | 4.67 ± 2.42 |  |

Note: M, mean; SD, standard deviation; CA, congenital amusia. Global coherence is presented as the number of individuals with / without global coherence. Informativeness efficiency is expressed in token words number per second. Informativeness density is resented as percentages. Local coherence is expressed in mean cosine similarity. Other metrics are expressed in counts.
\* *p* < 0.05; \*\* *p* < 0.01; \*\*\* *p* < 0.001

**Extended Data Table. 7.**
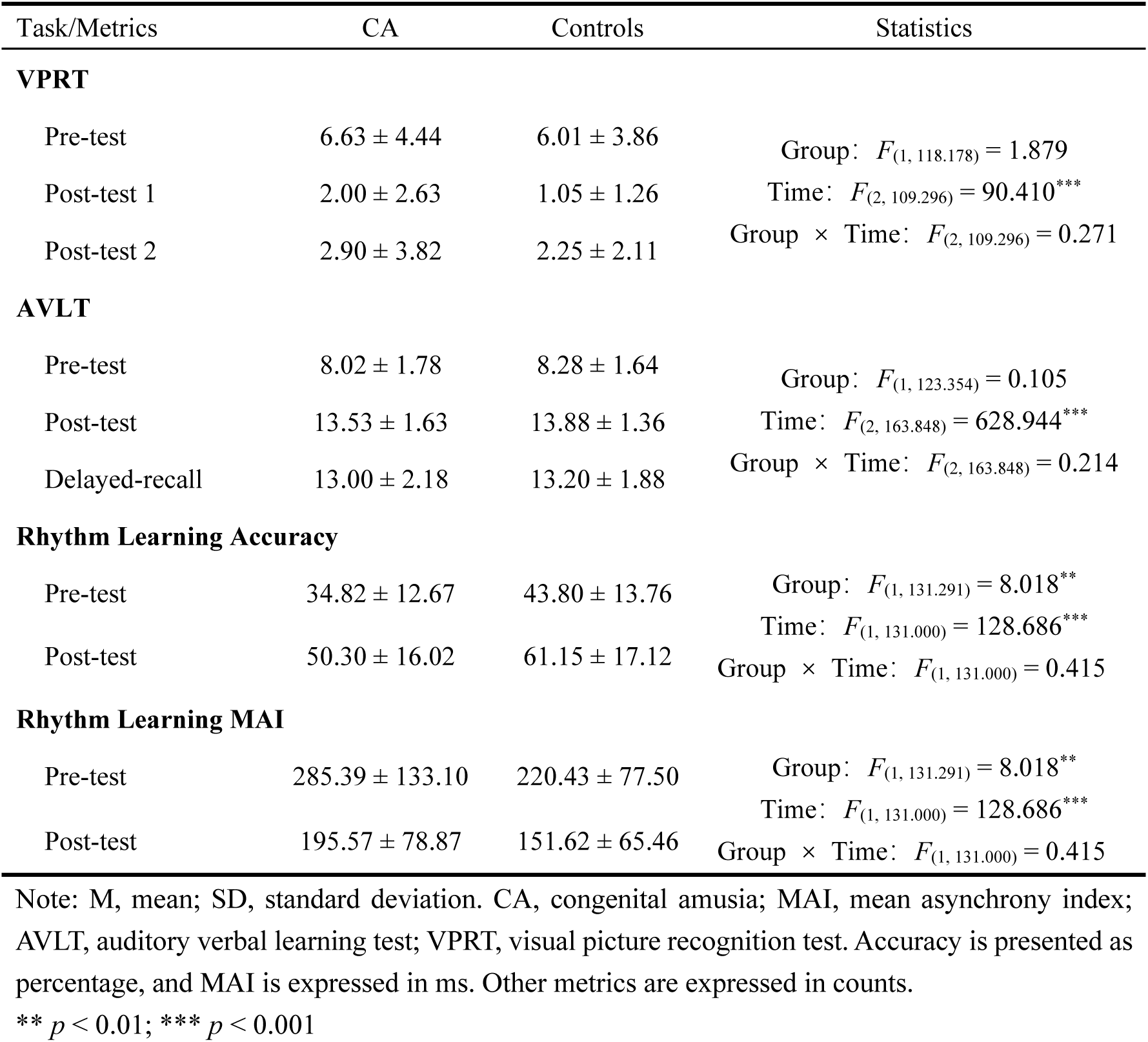
Episode memory and rhythm learning performance (M ± SD) and linear mixed model comparisons.

| Task/Metrics | CA | Controls | Statistics |
|---|---|---|---|
| **VPRT** |  |  |  |
| Pre-test | 6.63 ± 4.44 | 6.01 ± 3.86 | Group：*F*<sub>(1, 118.178)</sub> = 1.879<br>Time：*F*<sub>(2, 109.296)</sub> = 90.410<sup>\*\*\*</sup><br>Group × Time：*F*<sub>(2, 109.296)</sub> = 0.271 |
| Post-test 1 | 2.00 ± 2.63 | 1.05 ± 1.26 |  |
| Post-test 2 | 2.90 ± 3.82 | 2.25 ± 2.11 |  |
| **AVLT** |  |  |  |
| Pre-test | 8.02 ± 1.78 | 8.28 ± 1.64 | Group：*F*<sub>(1, 123.354)</sub> = 0.105<br>Time：*F*<sub>(2, 163.848)</sub> = 628.944<sup>\*\*\*</sup><br>Group × Time：*F*<sub>(2, 163.848)</sub> = 0.214 |
| Post-test | 13.53 ± 1.63 | 13.88 ± 1.36 |  |
| Delayed-recall | 13.00 ± 2.18 | 13.20 ± 1.88 |  |
| **Rhythm Learning Accuracy** |  |  |  |
| Pre-test | 34.82 ± 12.67 | 43.80 ± 13.76 | Group：*F*<sub>(1, 131.291)</sub> = 8.018<sup>\*\*</sup><br>Time：*F*<sub>(1, 131.000)</sub> = 128.686<sup>\*\*\*</sup><br>Group × Time：*F*<sub>(1, 131.000)</sub> = 0.415 |
| Post-test | 50.30 ± 16.02 | 61.15 ± 17.12 |  |
| **Rhythm Learning MAI** |  |  |  |
| Pre-test | 285.39 ± 133.10 | 220.43 ± 77.50 | Group：*F*<sub>(1, 131.291)</sub> = 8.018<sup>\*\*</sup><br>Time：*F*<sub>(1, 131.000)</sub> = 128.686<sup>\*\*\*</sup><br>Group × Time：*F*<sub>(1, 131.000)</sub> = 0.415 |
| Post-test | 195.57 ± 78.87 | 151.62 ± 65.46 |  |

Note: M, mean; SD, standard deviation. CA, congenital amusia; MAI, mean asynchrony index; AVLT, auditory verbal learning test; VPRT, visual picture recognition test. Accuracy is presented as percentage, and MAI is expressed in ms. Other metrics are expressed in counts.
\*\* *p* < 0.01; \*\*\* *p* < 0.001

