## Supplemental Information for "Neural competition predicts speech production advantages in congenital amusia"

### Supplementary Method 1. The INDI reference dataset

**INDI dataset selection.** The reference sample was drawn from INDI with the following inclusion criteria: (1) contributed by a Chinese research team, to reduce potential effects of ethnic background on imaging features, given that the our sample was entirely Chinese and INDI does not uniformly release participant ethnicity; (2) provision of rs-fMRI, sMRI and dMRI in the same participants together; (3) at least 30 diffusion-sensitive gradient directions, sufficient for probabilistic tractography; (4) inclusion of healthy participants; and (5) no dataset that was a universe or subset of another included dataset. On these criteria, data were obtained from the Beijing Enhanced dataset (https://fcon_1000.projects.nitrc.org/indi/retro/BeijingEnhanced.html), BNU 1, BNU 3, HNU 1, IPCAS 1, IPCAS 2, IPCAS 8 and XHCUMS sub-datasets of the CoRR datasets (https://fcon_1000.projects.nitrc.org/indi/CoRR/html/index.html), the SLIM dataset (https://fcon_1000.projects.nitrc.org/indi/retro/southwestuni_qiu_index.html) and the Beijing Short TR Study datasets (https://fcon_1000.projects.nitrc.org/indi/pro/BeijingShortTR.html).

Supplementary Table.1 Descriptive analysis of INDI participants' basic information and comparisons across datasets（M ± SD）

| Dataset | Sex（Male/Female） | Age（years） | Mean FD | IQR | TIV |
|---|---|---|---|---|---|
| Beijing Enhanced | 73/107 | 21.22 ± 1.94 | 0.06 ± 0.03 | 2.37 ± 0.17 | 1455.29 ± 130.53 |
| CoRR | | | | | |
| BNU 1 | 30/27 | 23.05 ± 2.29 | 0.06 ± 0.02 | 2.30 ± 0.11 | 1480.94 ± 129.15 |
| BNU 3 | 24/24 | 22.54 ± 2.15 | 0.06 ± 0.04 | 2.38 ± 0.14 | 1492.25 ± 121.88 |
| HNU 1 | 15/15 | 24.37 ± 2.41 | 0.04 ± 0.02 | 2.21 ± 0.34 | 1552.39 ± 138.83 |
| IPCAS 1 | 9/21 | 20.90 ± 1.77 | 0.07 ± 0.03 | 2.47 ± 0.28 | 1453.56 ± 130.71 |
| IPCAS 2 | 12/22 | 13.29 ± 1.00 | 0.07 ± 0.02 | 2.14 ± 0.10 | 1449.14 ± 101.19 |
| IPCAS 8 | 6/7 | 57.62 ± 3.78 | 0.10 ± 0.06 | 2.31 ± 0.16 | 1398.98 ± 155.94 |
| XHCUMS | 13/10 | 51.87 ± 6.41 | 0.08 ± 0.05 | 2.28 ± 0.10 | 1456.77 ± 186.43 |
| SLIM | 257/323 | 20.08 ± 1.28 | 0.06 ± 0.03 | 2.26 ± 0.14 | 1479.46 ± 127.28 |
| Beijing Short TR | 14/14 | 24.50 ± 7.52 | 0.07 ± 0.03 | 2.34 ± 0.13 | 1446.27 ± 132.30 |
| $\chi^2$/F | 9.037 | 945.560 | 5.925 | 23.915 | 2.767 |
| *p* | 0.434 | < 0.001 | < 0.001 | < 0.001 | 0.003 |

Note：Sex and age are reported based on all 1,023 participants; MRI-related variables are reported based on all participants finally included in the analysis for the corresponding modality. M: Mean; SD: standard deviation; FD: Frame-wise Displacement; IQR: Image Quality Rating; INDI: International Neuroimaging Data-sharing Initiative; CoRR: Consortium for Reliability and Reproducibility; BNU: Beijing Normal University; HNU: Hangzhou Normal University; IPCAS: Institute of Psychology, Chinese Academy of Sciences; XHCUMS: Xuanwu Hospital, Capital Medical University; SLIM: Southwest University Longitudinal Multimodal Brain Imaging Dataset.

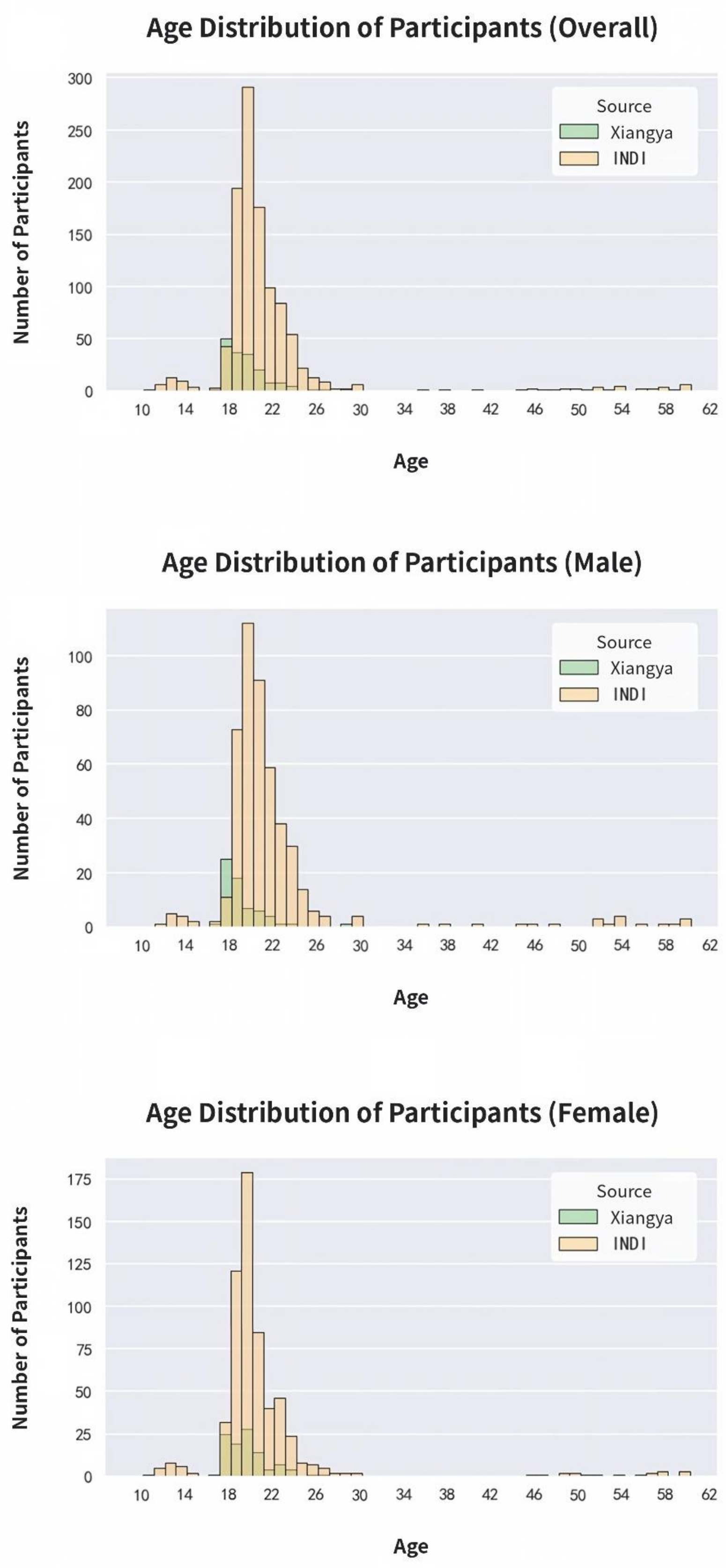


**Supplementary Fig.1**, Age distributions of INDI (apricot) and Xiangya samples (green) stratified by sex. From top to bottom, the panels show the age distributions for all participants, males and females, respectively. INDI, International Neuroimaging Data-sharing Initiative.

**Session selection for repeated scans.** CoRR was designed to assess the reproducibility of imaging metrics[1], and the same participant may therefore have multiple sessions. For each modality, only the earliest session in which that modality was acquired was retained. The same rule was applied to SLIM, which also contains repeated scans. Because the Beijing Short TR Study was designed to compare the effect of long and short repetition times on neuroimaging measures, and all other datasets used long TR for rs-fMRI, only long-TR rs-fMRI data from this dataset were retained to avoid confounding.

### Supplementary Method 2. Self-perceived Narrative Expression

To investigate participants' self-perceived narrative expression ability, an 8-item questionnaire was designed to collect relevant data. The items were as follows: (1) I can use appropriate language and vocabulary to accurately convey my thoughts and feelings during oral narrative expression; (2) I can narrate a complete story or experience clearly and coherently during oral narrative expression; (3) I can maintain logicality and organization during oral narrative expression, making it easier for listeners to understand; (4) I can remain calm and confident during oral narrative expression; (5 My oral narrative expression is expressive and engaging; (6) Most people familiar with me consider me good at oral narrative expression; (7) I am only willing to express myself in familiar settings and among familiar people; (8) I am unwilling to express myself even when necessary. All items were rated on a 5-point scale (items 7 and 8 were reverse-scored), with 1 indicating "strongly disagree" and 5 indicating "strongly agree"; higher scores indicated better self-perceived expressive ability.

### Supplementary Method 3. Image acquisition of Xiangya Samples and Image Data Processing

Scanning of Xiangya samples was performed at the Department of Radiology, the Second Xiangya Hospital, on a 3 T Siemens Skyra scanner. Before the research sequences, T2-weighted imaging (supplemented by sMRI and/or dMRI where clinically indicated if need) was used to screen for organic pathology; participants with incidental findings of clinical significance were excluded and referred.

rs-fMRI data were acquired using gradient-echo planar imaging sequence with following parameters: TR = 2,000 ms, TE = 30 ms, flip angle = 80°, 32 slices, matrix 256 × 256 $mm^2$, voxel size 4 × 4 × 4 $mm^3$, 216 volumes. rs-fMRI data were preprocessed with DPABI[2], including removal of the first 10 volumes, slice-timing correction, head-motion correction, normalization to standard Montreal Neurological Institute space (3 × 3 × 3 $mm^3$), smoothing (6 mm full-width at half-maximum kernel), regression of nuisance covariates (linear drift, white matter, cerebrospinal fluid, and motion parameters), and band-pass filtering (0.01–0.1 Hz). Four individuals with congenital amusia (CA), three controls and seven INDI participants were excluded from further analysis due to excessive head motion (mean frame-wise displacement > 0.2 mm). Finally, 65 individuals with CA and 94 controls in Xiangya dataset, and 1012 INDI participants entered the formal analysis. After preprocessing, we computed the metrics of interest. The amplitude of low-frequency fluctuations (ALFF)[3] is the mean amplitude within a specific frequency domain (here, 0.01–0.1 Hz) derived from a fast Fourier transform of a voxel's time course. For static ALFF (sALFF), values were first computed at the voxel level and converted to z-scores, and then averaged across all voxels within each IFG and STG subregion to yield ROI-level sALFF. Static functional connectivity (sFC) was computed as the Pearson correlation between the mean time series of each IFG subregion and that of each ipsilateral STG subregion and its contralateral homologue, followed by Fisher z transformation. DynamicBC[4] was used with a sliding-window approach to calculate dynamic metrics. The original time series was segmented into windows of 30 volumes with 80% overlap. Within each window, ALFF (voxel level) and FC were computed in the same way as for the static metrics. The variance of ALFF and FC across windows constituted dALFF and dFC, respectively; ROI-level dALFF was generated in the same way as sALFF. Because dynamic metrics are sensitive to sliding-window setting, all analyses were repeated with a 50-volume window and the same overlap to assess robustness.

sMRI data were acquired using magnetization-prepared rapid acquisition gradient-echo sequence with the following parameters: TR = 1,900 ms, TE = 2.03 ms, flip angle = 9°, 176 slices, acquisition matrix 256 × 256 $mm^2$, voxel size 1 × 1 × 1 $mm^3$. sMRI data were processed with CAT12, including origin correction, tissue segmentation, normalization, modulation and smoothing smooth (12 mm full-width at half-maximum kernel). After processing, GMV was extracted as the mean across all voxels within each ROI, and cortical thickness as the mean across all vertices within each ROI (we mapped ROI in voxel space to vertex space), from the modulated GMV and cortical thickness maps respectively. Two individuals with CA, six controls and thirty INDI participants were excluded from further analysis due to poor image quality (IQR > mean IQR + 2SD). Finally, 67 individuals with CA and 91 controls in

Xiangya dataset, and 986 INDI participants entered the formal analysis.

dMRI data were acquired using single-shot spin-echo planar imaging sequence with following parameters: TR = 7,000 ms, TE = 86 ms, flip angle = 90°, 60 slices, matrix 128 × 128 $mm^2$, voxel size 2 × 2 × 2.5 $mm^3$, 64 diffusion directions, b = 1,000 $s/mm^2$. dMRI data were preprocessed with FSL, including motion and eddy-current correction, gradient reorientation, brain extraction, and diffusion tensor fitting. We then conducted probabilistic tractography to reconstruct the arcuate fasciculus (AF). BedpostX was run with default parameters using Markov chain Monte Carlo sampling to estimate the distribution of diffusion parameters. ProbtrackX2 was then run with default parameters (5,000 samples per voxel), sampling repeatedly from the principal diffusion directions to build a prior distribution and infer the true fiber orientation. Tracking was performed bidirectionally between the two endpoint seed regions of each segment (Supplementary Fig.4a). During reconstruction, exclusion masks were applied to reduce spurious streamlines. These included the seed regions not traversed by each segment, the internal and external capsules, the corpus callosum, and voxels with FA < 0.1 (internal/external capsules and the corpus callosum were defined according to the Harvard-Oxford Atlas). In addition, two segment-specific exclusion masks were used. For the anterior segment, an exclusion mask was placed at the junction of the posterior temporal lobe and the inferior parietal lobule to exclude streamlines extending into the temporal lobe (Supplementary Fig.4b). For the posterior segment, an exclusion mask was placed along a plane extending from the anterior superior precentral gyrus to the posterior inferior postcentral gyrus to exclude streamlines extending into the frontal lobe (Supplementary Fig.4c). A connection-probability threshold of 0.1% was applied, calculated as the number of streamlines passing through a specific voxel divided by the total number of streamlines reaching the target, and voxels below this threshold were excluded. The union of the two directional reconstructions constituted the final tract. All reconstructions were visually inspected; failed reconstructions of the right direct segment occurred in two Xiangya controls and three INDI participants and were excluded from analyses involving that tract. All other tracts were successfully reconstructed in all participants. Based on the reconstructed arcuate fasciculus, the following metrics were calculated for each subsegment: (1) volume, expressed as the number of voxels occupied by the tract divided by the total number of whole-brain voxels; (2) fractional anisotropy (FA)/mean diffusivity (MD), the mean FA/MD value across all voxels of the tract; (3) axial diffusivity (AD), the mean of the first eigenvalues across all voxels of the tract; (4) radial diffusivity (RD), the mean of the average of the second and third eigenvalues across all voxels of the tract; (5) tract connection density, expressed as:

$$\frac{W_{AB}/V_B + W_{BA}/V_A}{2}$$

where $W_{AB}$ represents the number of streamlines originating from brain region A that were not excluded by the exclusion masks and ultimately reached brain region B, and $V_B$ represents the volume of brain region B; similarly, $W_{BA}$ represents the number of streamlines originating from brain region B that were not excluded and ultimately reached brain region A, and $V_A$ represents the volume of brain region A.

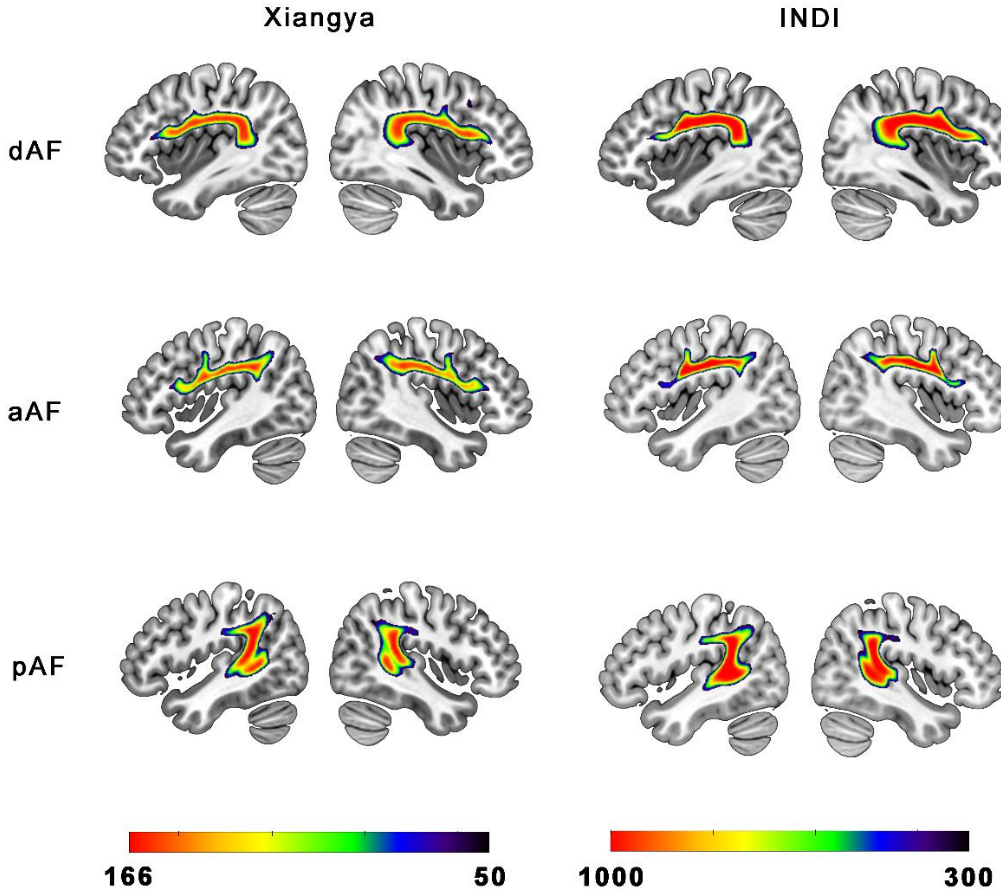


**Supplementary Fig.2 Reconstruction results of the arcuate fasciculus sub-segments in participants from the Xiangya and INDI.** For each panel, the left and right columns display the left and right hemispheres, respectively. The maps show common white matter tracts present in at least 30% of individuals; regions in red indicate areas identified as the corresponding arcuate fasciculus sub-segment in a greater proportion of individuals. The dAF, aAF and pAF are shown at x = ±38, x = ±42, and x = ±44, respectively. dAF, direct arcuate fasciculus; aAF, anterior arcuate fasciculus; pAF, posterior arcuate fasciculus; INDI, International Neuroimaging Data-sharing Initiative;

### Supplementary Method 4. ROI Definition

Supplementary Table.2 Brain regions of the Brainnetome atlas involved in this study

| Gyrus | Anatomical and modified Cyto-architectonic descriptions | Label ID | |
|---|---|---|---|
| | | L | R |
| Middle Frontal Gyrus | Inferior Frontal Junction [a] | 17 | 18 |
| | Dorsal Area 44 [a] | 29 | 30 |
| Inferior Frontal Gyrus | Inferior Frontal Sulcus | 31 | 32 |
| | Caudal Area 45 [a] | 33 | 34 |
| | Rostral Area 45 | 35 | 36 |
| | Opercular Area 44 | 37 | 38 |
| | Ventral Area 44 [a] | 39 | 40 |
| Precentral Gyrus | Caudal Ventrolateral Area 6 [a] | 63 | 64 |
| | Medial Area 38 | 69 | 70 |
| Superior Temporal Gyrus | Area 41/42 [b] | 71 | 72 |
| | TE 1.0/1.2 | 73 | 74 |
| | Caudal Area 22 [b] | 75 | 76 |
| | Lateral Area 38 | 77 | 78 |
| | Rostral Area 22 | 79 | 80 |
| posterior Superior Temporal Sulcus | Rostral [b] | 121 | 122 |
| | Caudal [b] | 123 | 124 |
| | Rostrodorsal Area 39 [c] | 137 | 138 |
| Inferior Parietal Lobule | Rostrodorsal Area 40 [c] | 139 | 140 |
| | Caudal Area 40 [c] | 141 | 142 |

Note: Superscripts a, b, and c denote the frontal, temporal, and parietal seed regions used for probabilistic tractography, respectively.

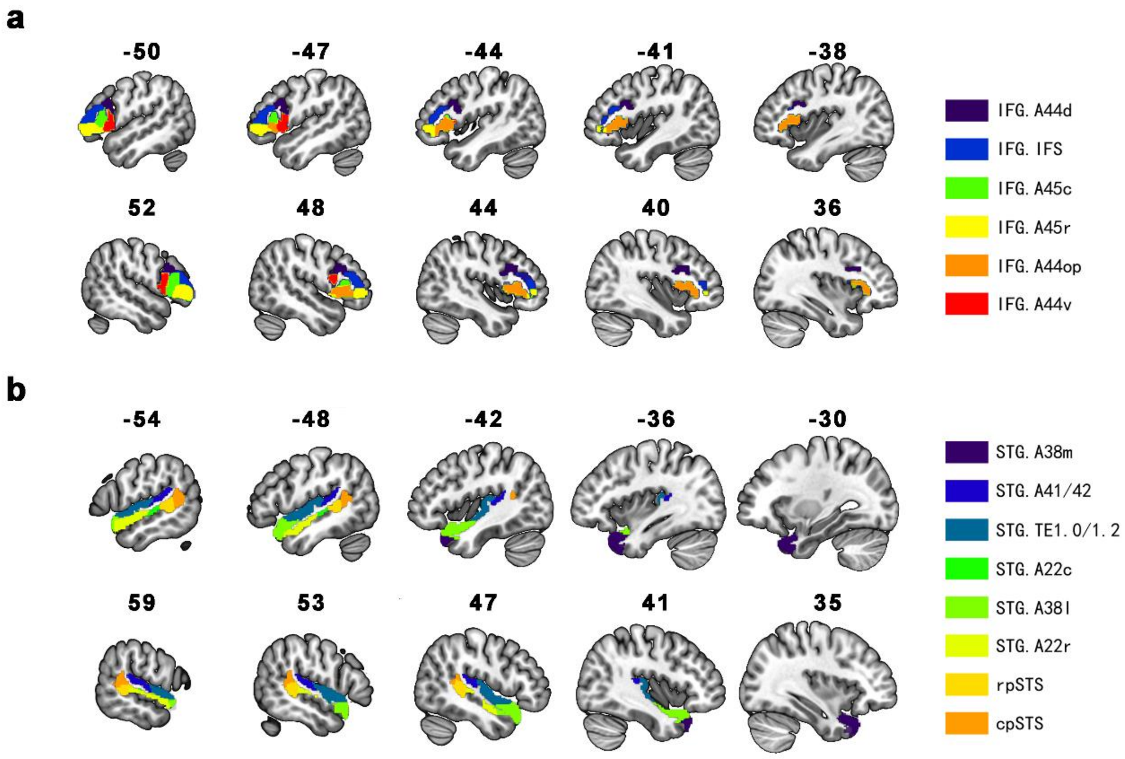


**Supplementary Fig.3 Sagittal anatomical locations of the regions of interest extracted from the Brainnetome atlas in the present study. a,** sub-regions of IFG. **b,** sub-regions of STG. Numbers on the brain maps indicate the x-axis coordinates of the corresponding slices in MNI space, with negative values indicating the left hemisphere and positive values indicating the right hemisphere. IFG, inferior frontal gyrus; STG, superior temporal gyrus; A44d, dorsal area of 44; A45c, caudal area of 45; IFS, inferior frontal sulcus; A45r, rostral area of 45; A44op, opercular area of 44; A44v, ventral area of 44; A38m, medial area of 38; A41/42, area of 41/42; A22c, caudal area of 22; A38l, lateral area of 38; A22r, rostral area of 22; rpSTS, rostral posterior superior temporal sulcus; cpSTS, caudal posterior superior temporal sulcus.

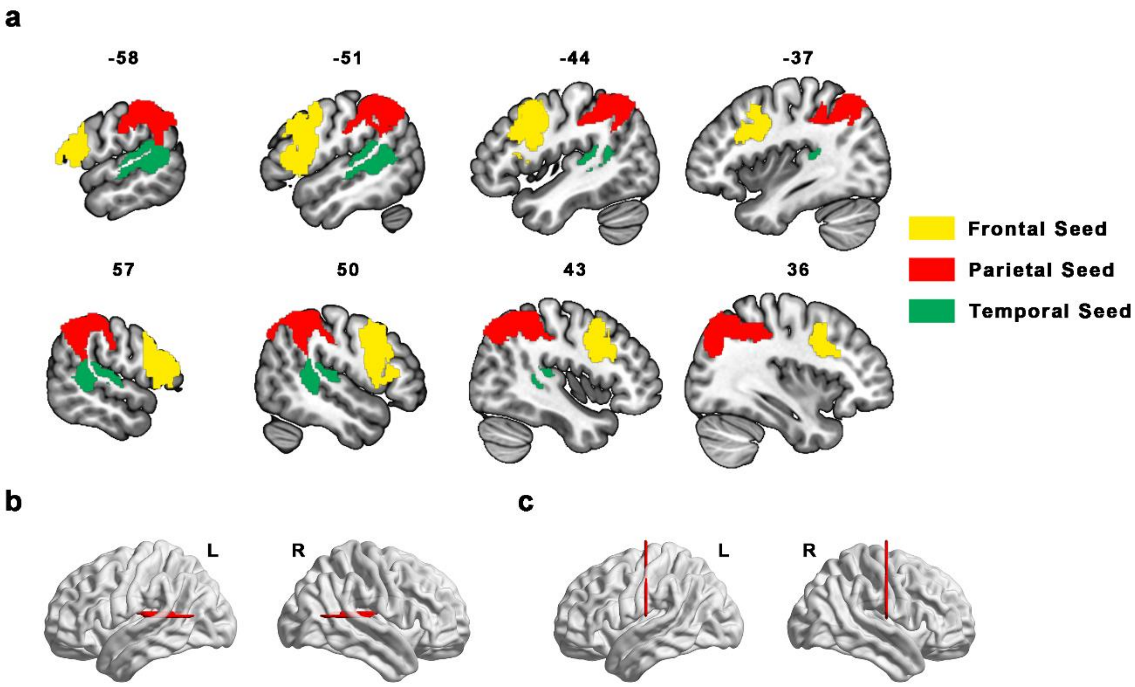


**Supplementary Fig.4 Anatomical locations of the masks for probabilistic tractography in the present study. a,** seed masks for the sub-segments of arcuate fasciculus; numbers on the brain maps indicate the x-axis coordinates of the corresponding slices in MNI space, with negative values indicating the left hemisphere and positive values indicating the right hemisphere. **b,** additional exclusion masks for the anterior arcuate fasciculus. **c,** additional exclusion masks for the posterior arcuate fasciculus. L, left; R, right.

### Supplementary Method 5. Metrics of Speech Production Task

#### Fluency

*Speech rate* was defined as the total number of syllables produced by the participant per unit time[5], reflecting the overall tempo and fluency of speech production.

*Silent pauses* were defined as silent intervals greater than 150 ms between adjacent syllables. Pauses were classified as short (150–400 ms), medium (400–1,000 ms) or long (>1,000 ms)[6]. For each category and for all silent pauses combined, frequency and duration proportion were computed.

Non-silent pauses, or termed *voiced pauses*, comprise *filled pauses*, *repetitions* and *repairs*[7]. *Filled pauses* were defined as non-intentional vocal or verbal fillers that occupy the speech stream without conveying planned linguistic content, such as “em” and “uh”. *Repetitions* were defined as immediate repetition of a word, character, syllable, sound, phrase or sentence by the speaker, where the repetition is semantically and pragmatically unnecessary. *Repairs* were defined as the speaker beginning an utterance, perceiving it as erroneous, inappropriate, unclear or insufficient, and then interrupting, abandoning or replacing it with another expression. Filled pauses, non-filled paused (i.e., repetitions and repairs) were identified from the original transcript and their frequencies (times per minute) were computed. Filled pauses and non-filled paused were combined as total voiced pause, and its frequencies (times per minute) were also computed.

*Total pause frequency* was computed as the number of silent plus voiced pauses per minute of narration.

**Micro-linguistic measures**

*Total number of words (TNW)* was the number of non-punctuation lexical tokens across all utterances, reflecting narrative productivity.

*Lexical diversity* was the number of different words among all words in the participant's narrative text, reflecting vocabulary reserve, flexibility in word use and the richness of verbal output during narration.

*Total number of utterances (TNU)* was the number of sentences obtained by segmenting the corrected transcript according to semantic and prosodic boundaries, reflecting narrative productivity.

*Mean length of utterance (MLU)* was total number of words divided by total number of utterances, reflecting the complexity of sentence construction.

*Syntactic complexity* was the proportion of utterances containing at least one complex syntactic structure[8], reflecting the complexity of sentence construction and the ability to organize complex syntax. Six structures were targeted, following previous work on Chinese[8]: *ba* constructions, *bei* constructions, pivotal constructions, serial-verb constructions, relative clauses and complement clauses. Structures were identified from part-of-speech tags, semantic role labels and dependency relations obtained during parsing.

**Macro-linguistic measures**

*Theme search* indexed identification, tracking and presentation of the narrative cues[9,10]. Four core themes were defined: awareness of being alone (theme 1, episode 1); desire to change the situation (theme 2, episode 2); meeting the elderly man and living together (theme 3, episodes 3–6); and return to isolation after the man's death (theme 4, episode 7). One point was awarded for each theme mentioned, giving a total score ranging from 0 to 4.

*Global coherence* indexed integration and presentation of the story's main storyline. Narration was judged as globally coherent if it accurately conveyed the main storyline, that is, a wandering orphan attempts to change his isolated state but ultimately returns to isolation, and as not globally coherent otherwise. A score of 1 was assigned if global coherence was present, and 0 if it was absent.

*Local coherence* indexed the semantic relatedness between adjacent utterances, reflecting the naturalness and fluency of meaning connection and transition between local utterances. Local coherence was quantified as the mean semantic relatedness strength between every pair of adjacent utterances.

*Narrative content* indexed extraction and presentation of narrative content. 30 panels of the were defined as thirty standard events; and core elements were specified for each event, including characters, actions, scene, interpersonal relations, character states and other necessary details. Each event was scored as accurate if all core elements were correctly described, as partially accurate if some elements were correct but incomplete, as error if any core element was described incorrectly, and as omitted if no content related to the event appeared in narration. Scores for the four categories each ranged from 0 to 30 and summed to 30.

*Informativeness density* was the proportion of token words among all produced words, indexing information density of the narrative. Token words are defined as accurate, intelligible, informative and content-relevant words.

*Informativeness efficiency* was the number of token words produced per minute, indexing the rate of information transmission.

### Supplementary Methods 6. Calculation for the Speech Metrics

**Transcription**. Recordings were transcribed with the large-model automatic speech recognition service provided by Volcengine (https://www.volcengine.com/docs/6561/1354868). Besides transcript, the service returned a text with utterance segmentation based on semantics and prosody, together with onset and offset times for each syllable. Transcript output in the stage is referred to as the original text. Narration duration could be represented as the interval between the onset of the first syllable and the offset of the last.

**Text correction**. HanLP (https://github.com/hankcs/HanLP)[11] was used to remove redundant text associated with disfluencies and to correct spelling and punctuation errors introduced during transcription, yielding the corrected text. Correction was restricted to disfluency-related redundancy and transcription errors; lexical and syntactic choices were preserved.

**Speech rate.** Speech rate was computed as the total number of syllables divided by total narration duration, where syllables were non-punctuation characters in the original text and duration was the interval between the onset of the first syllable and the offset of the last.

**Silent pause indices.** Frequency was computed as the number of silent pauses divided by narration duration, and silent pause duration proportion was computed as cumulative pause duration divided by narration duration. Silent pauses were identified based on the onset time of the *i-th* syllable and offset time of the *(i-1)-th* syllable obtained during transcription.

**Micro-linguistic indices.** LTP (https://github.com/HIT-SCIR/ltp)[12] was used for sentence segmentation, word segmentation, part-of-speech tagging, semantic role labelling and dependency parsing, as it outperforms HanLP on these tasks but lacks text-correction capability. TNU was the number of sentences produced by LTP. TNW was the number of non-punctuation words across all sentences. MLU was computed as TNW divided by TNU. Six complex structures were identified according to their structural rules using part-of-speech, semantic role and dependency information Syntactic complexity was the proportion of utterances containing at least one complex structure. Lexical diversity was computed using the moving average type–token ratio (MATTR) algorithm[13] with a 100-word window and a 1-word step across the punctuation-stripped text; within each window, the number of distinct words was divided by the window length, and values were averaged across all windows

**Local coherence.** Embedding was used to computed local coherence, which convert sentences into vectors that encode semantic information. The strength of semantic relatedness between utterances can be represented by the similarity between semantic vectors[14]. We used the text-embedding-v4 model[15] provided by Alibaba Cloud (https://help.aliyun.com/zh/model-studio/embedding) to convert each utterance in the corrected text (segmented as in the narrative utterance count) into a 1024-dimensional dense vector. The cosine similarity between the semantic vectors of adjacent sentence pairs was then used to represent semantic relatedness strength and to compute the local coherence.

**Informativeness**. Following previous studies[16,17], the token words in this study comprise nouns, verbs, adjectives, adverbs, numerals, classifiers, onomatopoeia and conjunctions, and were constructed in four steps. First, two researchers independently listed all such words directly depicted in the pictures or related to the story themes. Then, discrepancies were discussed and resolved, and the lists were merged, with one representative word retained per semantic item. Third, both researchers generated synonyms as many as possible for each retained word. Last, words used in 15 narrative texts elicited with the same materials but not included in the formal study were extracted, and any new token words and their synonyms were added. The final list contained 1,339 items. Because LTP uses the 863 part-of-speech tag set, which is more fine-grained than token words categories, tags n, nd, nh, ni, nl, ns, nt and nz were merged into nouns, and tags a and b were merged into adjectives. Some idiomatic token words in the final list were classified by adjective, verb or adverb, whereas the 863-tag set in LTP labels all idioms uniformly as *idiom*. We therefore identified, among token words tagged as idiom, those corresponding to adjective-, verb-, or adverb-type entries in the final list and counted them separately. Informativeness density was calculated as the number of token words in the transcript divided by the TNW. Informativeness efficiency was calculated as the number of token words in the transcript divided by duration of narration.

**Other Indices.** Large language models (LLMs) possess powerful capabilities in information extraction, semantic understanding, and generation, and their performance on text analysis tasks has been widely validated[18-25]. In this study, LLM (specifically, DeepSeek-V3.2 in reasoning mode; https://api-docs.deepseek.com/news-/news251201) was used to analyze *voiced pauses*, *theme search*, *global coherenc*e and *narrative content*. Prompts comprised role setting, task definition, base evaluating rules (for example, “no subjective inference; judgments based only on explicit textual content”), measurement definitions, measurement-specific rules, workflow, few-shot examples, a JSON output schema and the text to be analyzed. Initial prompts were iteratively refined on 10 manually annotated texts not included in the formal study. Each iteration located discrepancies between model and human scores, examined the model's reasoning to identify their source, and distilled general rules that were integrated into the prompt (for example, adding "repetition inherent to conventional expressions is not counted as repetition" as measurement-specific rule and providing examples). Iteration continued until agreement was stable across two consecutive evaluations: a difference of $\leq 2$ in voiced pauses, exact agreement for each theme search and global coherence item, and Cohen's $\kappa \geq 0.9$ for event content. Generalization of the optimized prompts were tested on 5 additional manually annotated texts (elicited with the same materials but not included in the formal study). Results showed that (1) for voiced pauses, the difference between LLM and manual scores was ≤ 3 in four samples and 5 in one sample; (2) for theme search and global coherence, LLM outputs matched manual annotations exactly across all five samples; and (3) for narrative content, Cohen's κ between LLM and manual scores exceeded 0.8 in all five samples. These results indicate that the optimized prompts can accurately evaluate the target metrics. To guard against instability from insufficient learning of edge cases during prompt optimization and to reduce hallucination, a second LLM session independent with the main session analyzed the formal data in parallel using identical prompts. If the two sessions agreed, the main session's output was adopted; if not, the main session's output was manually reviewed and corrected by an operator blinded to participants' group and time-point labels. Agreement was defined as a difference of ≤ 3 in total voiced pauses, exact agreement on each theme search and global coherence item, and Cohen's κ ≥ 0.8 for narrative content.

### Supplementary Method 7. Example Pictures of Visual Picture Recognition Test

The task comprised three blocks, and the recognition phase of each block contained 50 pictures. The 50 pictures comprised 10 targets, 20 highly similar lures (HSLs) and 20 low similar lures (LSLs). HSLs are discriminable from the targets (Supplementary Fig.5a) only by the action depicted on the stimuli but not by its superficial characteristics, such as the presence or absence of people and objects (Supplementary Fig.5b); LSLs had no similarity with the targets, neither with regard to activity nor to surface characteristics (Supplementary Fig.5c). To avoid systematic bias caused by difficulty differences, the visual complexity of the pictures and the similarity between targets and lure pictures were counterbalanced across blocks. Each picture appeared only once in the entire test, and all pictures in each stage of the test were presented in a pseudo-random order. The task was programmed using JavaScript.

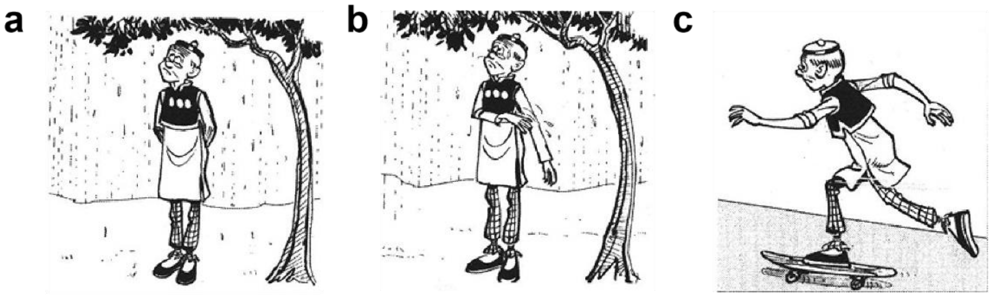


**Supplementary Fig.5 Examples of the three types of pictures in the recognition phase of the Visual Picture Recognition Test. a,** target. **b,** highly similar lure which is discriminable from the target only by the action depicted on the stimuli. **c,** low similar lure with no similarity to the targets.

### Supplementary Method 8. Rhythm Learning Task

The task comprised three blocks, each containing 12 trials. Each trial consisted of a listening phase and an imitation phase (Supplementary Fig.6a). During the listening phase, participants learned an 11-note rhythm sequence (lasted approximately 6 s) without using bodily movements to aid memorization. During the imitation phase, participants tapped the SPACE button of keyboard to synchronize with the rhythm as accurately as possible. The rhythms used in the three blocks were composed of the same 11 notes, but different permutations of note order formed three distinct rhythms. The 11-note rhythm comprised five eighth notes (the next note was played 50 ms after the end of the current note; the same convention applies to the descriptions of the other note durations below), three quarter notes (300 ms), one dotted quarter note (550 ms), one half note (800 ms), and one dotted half note (1300 ms). The duration of each note was 200 ms. A warning tone was presented 500 ms before each target sequence to signal the start of the trial. The task was programmed using JavaScript.

Task performance was evaluated by accuracy and mean asynchrony index (MAI). Accuracy was defined as the percentage of notes with correct responses relative to the total number of notes, with a correct response defined as a tap occurring within the correct response window. This window extended from after the midpoint of the preceding note's duration plus interval to before the midpoint of the current note's duration plus interval. Asynchrony was the absolute difference between tap time and note onset time, and MAI was the mean asynchrony over all correct responses. Supplementary Fig. 6b presents a schematic illustration of the calculation methods for the two indices.

Supplementary Table.2 Notes sequence of each block

| Block Number | Notes Sequence |
|---|---|
| 1 | 48842884̇842̇ |
| 2 | 48824̇8882̇44 |
| 3 | 284882̇44̇488 |

Note: 4 represents quarter note; 8 represents eighth note; 2 represents half note; 4̇ represents dotted quarter note and 2̇ represents dotted half note.

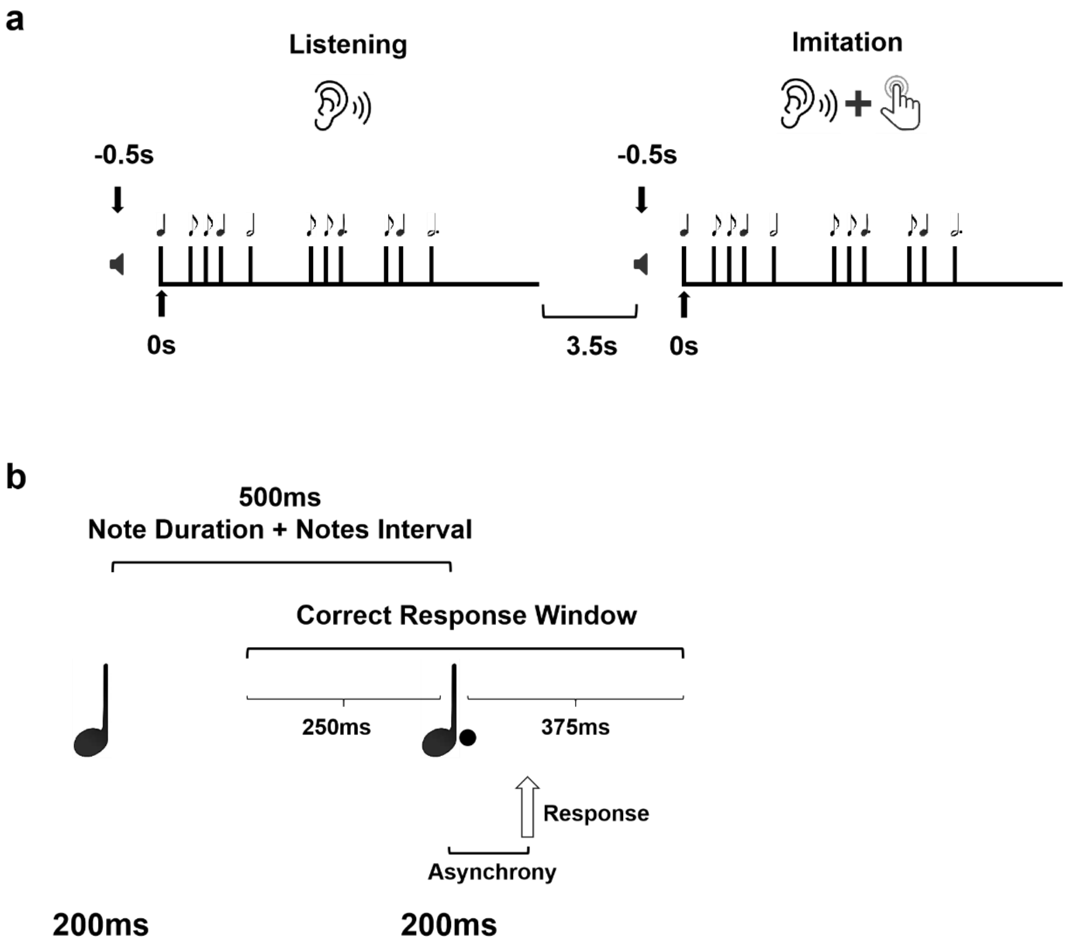


**Supplementary Fig.6 Illustration of the music rhythm learning task. a, a** complete trial consisted of two phases (listening and imitation), with a warning tone presented 500 ms before the rhythm sequence began**. b,** schematic illustration of the calculation methods for accuracy and mean asynchrony index. Each note had a duration of 200 ms, with inter-note intervals of 300 ms following quarter notes and 550 ms following dotted quarter notes. Therefore, for the dotted quarter note in the example, the correct response window spanned from 250 ms before the onset of the note to 375 ms after its onset. Asynchrony was calculated as the absolute difference between the tap time and the actual note onset time.

### Supplementary Discussion 1. Heterogeneity in the structural features of the right frontotemporal circuit in CA

Two findings in the structural features of the right frontal-temporal circuit (FTC) diverged from previous reports. First, unilateral right direct arcuate fasciculus (dAF) volume did not differ between groups in the present study, while dAF volume lateralization negatively deviated more often in CA (i.e., more subjects with relatively larger right dAF volume). Loui et al. found a reduction in the volume of dAF in CA[26], whereas other studies did not find differences in the volume of this structure between CA and controls[27,28]. Taken together with the present findings, these results suggest that CA may be characterized by heterogeneity in the volume of dAF. Specifically, this heterogeneity manifests as the coexistence of two opposite patterns of structural change: some individuals show a reduction in right-hemisphere arcuate fasciculus volume, whereas others show a relative increase in the volume of this structure. These two opposing structural patterns cancel each other out when averaged at the group level, thereby leading to the absence of significant group differences in the absolute volume of the unilateral arcuate fasciculus in the present study and in some previous studies. Combined with the present analyses of white matter microstructure and fiber connection density, we further speculate that this heterogeneity may reflect different neurodevelopmental trajectories associated with CA: (1) in some individuals, axonal demyelination or axonal loss arising from various causes is not followed by repair, resulting in impaired white matter microstructure and reduced volume[29-32]; (2) in some individuals, the brain may detect abnormal efficiency of neural signal propagation in this segment and initiate repair mechanisms, but the process does not proceed toward restoring normal white matter integrity and connection efficiency (e.g., axonal swelling)[33-35], ultimately producing a maladaptive state in which microstructural impairment coexists with increased volume; and (3) abnormal neural pruning during critical developmental periods may result in the retention of redundant neural connections between the right frontal and temporal lobes in some individuals, further disrupting the developmental trajectory of nerve fibers[36-39], ultimately leading to the coexistence of impaired white matter microstructure and increased macroscopic volume. Owing to differences in neurodevelopmental processes, when the proportions of individuals with different neurodevelopmental trajectories differ across studies (especially small-sample studies), inconsistent group-level conclusions may be reached. Through individual-level analysis, the present study successfully revealed neuro-phenotypic differences masked by group-average characteristics, which not only provides a new explanatory framework for reconciling the contradictions in previous studies but also offers a new perspective for understanding the heterogeneity of the neural basis of CA. Whether the different patterns of dAF volume change caused by different neurodevelopmental processes are associated with different cognitive and behavioral phenotypes of CA remains to be further elucidated in future studies.

Second, right IFG cortical thickness showed no group difference. This pattern is consistent with the findings of Liao et al.[40] but differs from those of Hyde et al.[41]. We suggest that the discrepancies among studies are mainly related to heterogeneity in the included participants. On the one hand, the ethnic background of participants may influence the results: the participants in the present study and in Liao et al.'s study were mainly East Asian, whereas those included in Hyde et al.'s study were mainly from European and American countries. Different ethnic groups may differ in cultural/genetic background, the structural basis of the brain, and the developmental trajectory of auditory cognitive functio[42-45]; these factors may directly affect the developmental process of cortical thickness in the FTC and its association patterns with CA, thereby leading studies based on different ethnic groups to obtain different results. On the other hand, sampling bias may likewise be an important factor contributing to inconsistent findings: owing to limited sample sizes and the possible existence of different behavioral subtypes of CA, the randomness of sampling may lead to systematic biases in the cortical thickness characteristics of the FTC among participants included in a given study, thereby affecting the stable detection of association patterns between cortical thickness alterations and CA. Therefore, future studies are needed in larger datasets covering different ethnic backgrounds, while incorporating potential subtypes of CA into the analytical framework, so as to more systematically and comprehensively elucidate the cortical thickness characteristics of the FTC in CA.

### Supplementary Discussion 2. Discrepancies across studies regarding left frontotemporal functional connectivity in CA

Although Jin et al. found weakened functional connectivity between the right inferior frontal gyrus (IFG) and the right superior temporal gyrus (STG) in CA, further supporting the view that functional blunting of right FTC is a neural basis of CA, that study also showed that functional connectivity between the left IFG and STG was likewise weakened in CA[46]. This is contrary to the findings and theoretical hypotheses of our study. Similar to the discussion of discrepancies across studies regarding the volume of dAF in CA, we suggest that this inconsistency may arise from differences in the specific manifestations of neural plasticity and from methodological differences. First, different neuroimaging phenotypes may be associated with several distinct potential manifestations of neural plasticity. First, owing to a decline in the functional efficiency of the right FTC, the homologous circuit in the left hemisphere may also undergo adaptive downregulation to maintain relative interhemispheric homeostasis of functional activity and ensure the normal operation of overall psychological function[47,48]. Second, the decline in functional efficiency of the right FTC may reduce the overall competitiveness of the FTC for neural resources in the brain, thereby leading to a synchronous weakening of functional connectivity in the homologous left-hemisphere circuit. However, the enhanced left-hemisphere functional connectivity observed in the present study strongly supports a third potential manifestation of neural plasticity, namely interhemispheric neural competition. The present study found that functional connectivity between the left IFG and the STG was negatively correlated with musical perception ability. This result indicates that the enhanced functional connectivity is not a compensation for the weakened functional connectivity between homologous brain regions in the right hemisphere, but rather arises because structural damage to key white matter pathways in the right FTC weakens the efficiency of functional coordination between the right IFG and STG and its inhibitory regulation of the left hemisphere, thereby releasing neural resources for or creating a “competitive vacuum” for the homologous left-hemisphere circuit, which in turn enhances its functional connectivity.

Methodologically, prior studies compared group means in small samples, whereas we combined group-level analysis in a relatively large sample with normative modelling. Different manifestations of neuroplasticity may produce heterogeneity of neuro-phenotypes. Small-sample group-mean analyses are poorly suited to capture such heterogeneity, and their conclusions may therefore vary depending on which phenotypes a given sample happens to overrepresent. Individual-level modelling can reveal deviations masked by group averaging. The discrepancy is therefore more plausibly due to sample size, metric choice and level of analysis than to a substantive inconsistency.

### Supplementary Discussion 3. Dissociation between subjective and objective measures, and alternative explanations for the behavioral advantages of speech production

Although CA demonstrated objective advantages in speech production, they did not subjectively report better narrative ability than controls. This dissociation may arise for two reasons. First, the advantages observed here, such as maintaining syntactic complexity and local coherence under load, arise from largely automatic and implicit processes whose outputs are not readily available to introspection. CA may therefore not recognize these behavioral features, so these advantages are unlikely to be integrated into self-awareness. Second, self-perceived speaking ability may be based mainly on performance in everyday speaking situations, such as appeal, clarity and persuasiveness. The advantages captured here are analytic psycholinguistic dimensions that individuals may not equate with being a good speaker. This mismatch between objective metrics and subjective conceptions of eloquence may prevent these advantages from translating into corresponding self-efficacy.

Two compensation accounts merit consideration. The first holds that CA may compensate by strengthening non-prosodic semantic output to ensure semantic accuracy, because impaired pitch production prevents them from using intonation and stress to convey meaning. This account does not fit the available evidence because intonation production in natural speech is largely preserved in CA[49-52], so there is no functional gap that would call for such compensation. The second holds that CA may develop verbalization strategies to encode musical information they cannot perceive accurately, and that these strategies generalize to everyday speech production. This account is also not supported by existing evidence, as CA tend to down-weight unreliable acoustic cues (e.g., pitch) and rely on other cues (e.g., duration) rather than verbalizing musical information[53,54]. Neither compensation account plausibly explains the present findings. The distinctive features CA showed in narrative speech production are therefore more likely to reflect advantages arising from neural competition, although this interpretation requires more direct empirical support.

7 Engelhardt, P. E., Nigg, J. T. & Ferreira, F. Is the fluency of language outputs related to individual differences in intelligence and executive function? *Acta*

*Psychol (Amst)* **144**, 424-432 (2013).
8 Torng, P.-C. & Sah, W.-H. Narrative abilities of Mandarin-speaking children with and without specific language impairment: macrostructure and microstructure. *Clinical Linguistics & Phonetics* **34**, 453-478 (2019).
9 Ash, S. *et al.* Trying to tell a tale: discourse impairments in progressive aphasia and frontotemporal dementia. *Neurology* **66**, 1405-1413 (2006).
10 Mueller, K. D., Hermann, B., Mecollari, J. & Turkstra, L. S. Connected speech and language in mild cognitive impairment and Alzheimer's disease: A review of picture description tasks. *Journal of Clinical and Experimental Neuropsychology* **40**, 917-939 (2018).
11 He, H. & Choi, J. D. 5555-5577 (Association for Computational Linguistics).
12 Che, W., Feng, Y., Qin, L. & Liu, T. 42-49 (Association for Computational Linguistics).
13 Covington, M. & McFall, J. Cutting the Gordian knot: The moving-average type-token ratio (MATTR). *Journal of Quantitative Linguistics* **17**, 94-100 (2010).
14 Seuffert, S. *et al.* Linking speech patterns to brain structure in affective and psychotic disorders: an integrative natural language processing approach. *Molecular Psychiatry* **31**, 2057-2068 (2025).
15 Zhang, Y. *et al.* Qwen3 Embedding: Advancing Text Embedding and Reranking Through Foundation Models. *ArXiv* (2025).
16 Lambon Ralph, M. A., Conroy, P., Halai, A. D. & Alyahya, R. S. W. A unified model of post-stroke language deficits including discourse production and their neural correlates. *Brain* **143**, 1541-1554 (2020).
17 Kim, H. & Kintz, S. Discourse Production Across the Adult Lifespan: Microlinguistic Processes. *Topics in Cognitive Science* (2024).
18 Hashir, M. H., Memoona & Kim, S. W. TARGE: large language model-powered explainable hate speech detection. *PeerJ Computer Science* **11**, e2911 (2025).
19 Rathje, S. *et al.* GPT is an effective tool for multilingual psychological text analysis. *Proceedings of the National Academy of Sciences* **121**, e2308950121 (2024).
20 Wiest, I. C. *et al.* Privacy-preserving large language models for structured medical information retrieval. *NPJ Digital Medicine* **7**, 257 (2024).
21 Zhu, H. *et al.* A systematic review and Bayesian network meta-analysis on the efficacy and potential of mobile interventions for stress management. *Nature Human Behaviour* **9**, 1431-1441 (2025).
22 Bermejo, V. J., Gago, A., Gálvez, R. H. & Harari, N. LLMs outperform outsourced human coders on complex textual analysis. *Scientific Reports* **15**, 40122 (2025).
23 Deiner, M. S. *et al.* Large Language Models Can Enable Inductive Thematic Analysis of a Social Media Corpus in a Single Prompt: Human Validation Study. *JMIR Infodemiology* **4**, e59641 (2024).
24 Davidson, T. Multimodal large language models can make context-sensitive hate speech evaluations aligned with human judgement. *Nature Human Behaviour* **10**, 514-530 (2025).
25 Wright, A. G. C. *et al.* Assessing personality using zero-shot generative AI

scoring of brief open-ended text. *Nature Human Behavior* **10**, 541-555 (2026).

26 Loui, P., Alsop, D. & Schlaug, G. Tone Deafness: A New Disconnection Syndrome? *Journal of Neuroscience* **29**, 10215-10220 (2009).

27 Chen, X. *et al.* The lateralized arcuate fasciculus in developmental pitch disorders among mandarin amusics: left for speech and right for music. *Brain Structure and Function* **223**, 2013–2024 (2018).

28 Chen, J. L. *et al.* Detection of the arcuate fasciculus in congenital amusia depends on the tractography algorithm. *Frontiers in Psychology* **6**, 9 (2015).

29 Zhang, S. *et al.* Mediator MED23 controls oligodendrogenesis and myelination by modulating Sp1/P300-directed gene programs. *Cell Discovery* **10**, 102 (2024).

30 Weier, K. *et al.* Impaired growth of the cerebellum in pediatric-onset acquired CNS demyelinating disease. *Mult Scler* **22**, 1266-1278 (2016).

31 Reilly, T. C. *et al.* A whole-brain structural connectivity signature in adult Niemann-Pick disease type C. *Brain Communications* **7**, fcaf426 (2025).

32 Rocca, M. A. *et al.* Clinically Isolated Syndrome Suggestive of Multiple Sclerosis: Dynamic Patterns of Gray and White Matter Changes—A 2-year MR Imaging Study. *Radiology* **278**, 841-853 (2016).

33 Maillard, A. M. *et al.* Pervasive alterations of intra-axonal volume and network organization in young children with a 16p11.2 deletion. *Translational Psychiatry* **14**, 95 (2024).

34 Ackerman, J. M., Ford, T. J. L., Kattewar, S. S. & Kim, W.-Y. Perineuronal nets in the developing brain: implications for neurodevelopmental disorders. *Molecular Brain* **18**, 88 (2025).

35 Paus, T., Keshavan, M. & Giedd, J. N. Why do many psychiatric disorders emerge during adolescence? *Nature Reviews Neuroscience* **9**, 947-957 (2008).

36 Schafer, Dorothy P. *et al.* Microglia Sculpt Postnatal Neural Circuits in an Activity and Complement-Dependent Manner. *Neuron* **74**, 691-705 (2012).

37 Liddelow, S. A. & Barres, B. A. Reactive Astrocytes: Production, Function, and Therapeutic Potential. *Immunity* **46**, 957-967 (2017).

38 Tang, G. *et al.* Loss of mTOR-dependent macroautophagy causes autistic-like synaptic pruning deficits. *Neuron* **83**, 1131-1143 (2014).

39 Mizuno, H. *et al.* NMDAR-Regulated Dynamics of Layer 4 Neuronal Dendrites during Thalamocortical Reorganization in Neonates. *Neuron* **82**, 365-379 (2014).

40 Liao, X., Sun, J., Jin, Z., Wu, D. & Liu, J. Cortical Morphological Changes in Congenital Amusia: Surface-Based Analyses. *Frontiers in Psychiatry* **12**, 721720 (2022).

41 Hyde, K. L. *et al.* Cortical thickness in congenital amusia: when less is better than more. *Journal of Neuroscience* **27**, 13028-13032 (2007).

42 Heckbert, S. R. *et al.* Association of quantitative susceptibility mapping signal in gray matter with clinical characteristics and aging-related MRI markers: The Multi-Ethnic Study of Atherosclerosis. *NeuroImage: Clinical* **48**, 103884 (2025).

43 Shimada, K. *et al.* Ethnic differences in COMT genetic effects on striatal grey matter alterations associated with childhood ADHD: A voxel-based morphometry study in a Japanese sample. *World Journal of Biological Psychiatry* **18**, 322-328 (2017).

44 Jednoróg, K. *et al.* How reliable are gray matter disruptions in specific reading disability across multiple countries and languages? Insights from a large-scale voxel-based morphometry study. *Human Brain Mapping* **36**, 1741-1754 (2015).

45 Jorgenson, E. *et al.* Ethnicity and Human Genetic Linkage Maps. *The American Journal of Human Genetics* **76**, 276-290 (2005).

46 Jin, Z. *et al.* Altered resting-state connectivity of the auditory cortex in congenital amusia: A functional magnetic resonance imaging study in Mandarin speakers. *Annals of the New York Academy of Sciences* **1541**, 140-150 (2024).

47 Graziadio, S., Tomasevic, L., Assenza, G., Tecchio, F. & Eyre, J. A. The myth of the 'unaffected' side after unilateral stroke: Is reorganisation of the non-infarcted corticospinal system to re-establish balance the price for recovery? *Experimental Neurology* **238**, 168-175 (2012).

48 Zappasodi, F. *et al.* Longitudinal quantitative electroencephalographic study in mono-hemispheric stroke patients. *Neural Regeneration Research* **14**, 1237-1246 (2019).

49 Liu, F. *et al.* Pitch perception and production in congenital amusia: Evidence from Cantonese speakers. *Journal of the Acoustical Society of America* **140**, 563 (2016).

50 Nan, Y., Sun, Y. & Peretz, I. Congenital amusia in speakers of a tone language: association with lexical tone agnosia. *Brain* **133**, 2635-2642 (2010).

51 Yang, W. X., Feng, J., Huang, W. T., Zhang, C. X. & Nan, Y. Perceptual pitch deficits coexist with pitch production difficulties in music but not Mandarin speech. *Frontiers in Psychology* **4**, 1024 (2013).

52 Zhu, J., Xu, M. & Shao, J. Individuals with congenital amusia show degraded performance in a nonword repetition task with lexical tones. *Cortex* **196**, 140-154 (2026).

53 Jasmin, K., Dick, F., Holt, L. L. & Tierney, A. Tailored perception: Individuals' speech and music perception strategies fit their perceptual abilities. *Journal of Experimental Psychology: General* **149**, 914-934 (2020).

54 Jasmin, K., Dick, F., Stewart, L. & Tierney, A. T. Altered functional connectivity during speech perception in congenital amusia. *eLife* **9**, e53539 (2020).
